# Robust Production of Uniform Human Cerebral Organoids from Pluripotent Stem Cells

**DOI:** 10.1101/2020.03.08.979013

**Authors:** A. Sivitilli, J.T. Gosio, B. Ghoshal, A. Evstratova, D. Trcka, P. Ghiasiighorveh, J. J. Hernandez, J. M. Beaulieu, J. L. Wrana, L. Attisano

**Author notes:** Corresponding author: Liliana Attisano, Donnelly Centre, Rm 1008, University of Toronto, 160 College Street, Toronto, ON, M5S 3E1, Canada.

## Abstract

Human cerebral organoid (hCO) models offer the opportunity to understand fundamental processes underlying human specific cortical development and pathophysiology in an experimentally tractable system. While diverse methods to generate brain organoids have been developed, a major challenge has been the production of organoids with reproducible cell type heterogeneity and macroscopic morphology. Here, we directly addressed this problem by establishing a robust production pipeline to generate morphologically consistent (ie uniform) hCOs and achieve a success rate of >80%. These hCOs include both a radial glial stem cell compartment and electrophysiologically competent mature neurons. Moreover, we show using immunofluorescence microscopy and single cell profiling, that individual organoids display reproducible cell type compositions that are conserved upon extended culture. We expect that application of this method will provide new insights into brain development and disease processes.

## Introduction

The development, patterning, and homeostatic maintenance of the human brain is complex and while considerable insights into mechanisms driving these processes have been obtained from studies in model organisms, species specific differences in brain development and function can make it challenging to apply results from animal models to humans. Accordingly, understanding the molecular basis underlying normal development, disease progression, and therapeutic options for human brain-associated diseases, including cancer, requires human models.

The ability to generate brain organoids derived from human pluripotent stem cells provides an unprecedented opportunity to study context-dependent human disease pathologies in an experimentally tractable system. Indeed, this approach has provided insights into alterations associated with Alzheimer’s, blindness, ASD, Zika virus infection and others (Amin & Pasca, 2018, Chen, Song et al., 2019, Di Lullo & Kriegstein, 2017, Lancaster & Knoblich, 2014b, Quadrato, Brown et al., 2016, Rossi, Manfrin et al., 2018). A variety of protocols to generate brain organoids have been developed, but the considerable variability and heterogeneity between individual organoids obtained using these methods limits the utility of the model for studying disease mechanisms, or for examining the therapeutic potential of new drug candidates. Here, we establish a robust protocol to efficiently and reproducibly generate mature, consistent human Cerebral Organoids (hCOs). By optimizing an established protocol for self-patterned whole-brain organoids (Lancaster & Knoblich, 2014a, Lancaster, Renner et al., 2013), we generated phenotypically consistent forebrain organoids with reproducible morphologies and cell-type compositions. Thus, this protocol is ideally suited for studying mechanisms underlying human diseases and for investigation of potential novel therapeutic options in an experimentally tractable system.

## Results

### Optimization of cerebral organoid production

To establish a method to produce highly similar brain organoids (**Fig. 1a**), we explored modifications to a previously established protocol for generating self-patterned whole-brain organoids (Lancaster & Knoblich, 2014a, Lancaster et al., 2013), which yields organoids with variable morphology and cell type composition (Quadrato, Nguyen et al., 2017, Velasco, Kedaigle et al., 2019, Yoon, Elahi et al., 2019). We primarily used female H9 human Embryonic Stem Cells (hESCs), and also validated results in a male hESC model (H1; see below). We first optimized embryoid body (EB) generation by plating singularized H9 cells into 96-well plates with variable geometries and surface coatings, and examined EBs of uniform shape at 5 days. In contrast to the irregular clusters observed in traditional U-bottom dishes with or without surface coatings, EB aggregates that formed in plates with non-adherent surfaces in a V bottom or Aggrewell™ geometry formed single, similarly sized spheres of 400-450 μm diameter in each V shaped well, all of which displayed similar opacity under bright field microscopy (**Fig. 1b-d**). As multiple EBs are generated using the Aggrewell system, it was technically challenging to isolate intact individual spheroids for differentiation, so we focused on the single “V” bottom non-binding format for all subsequent studies.

**Figure 1.**
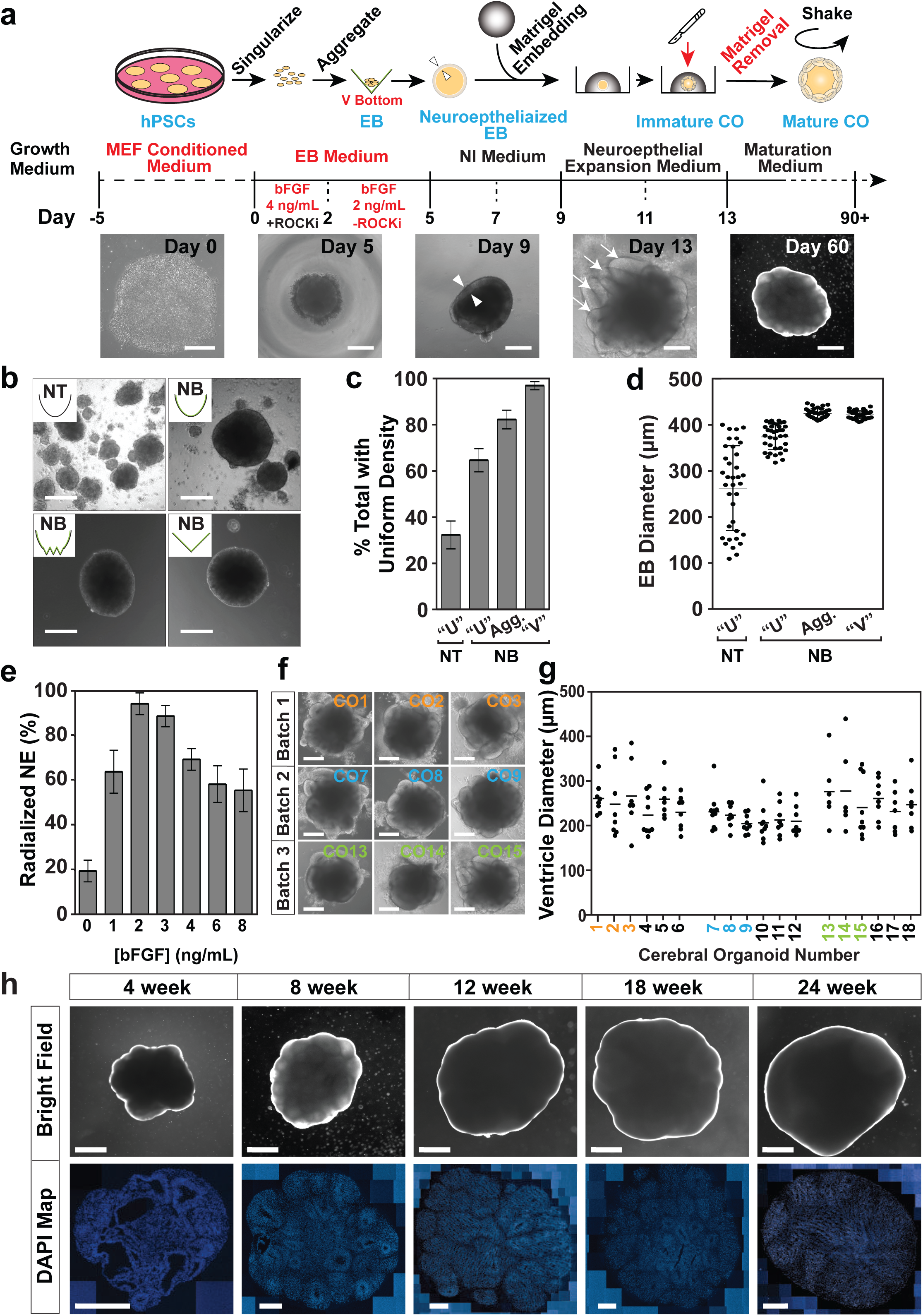
Generation of hCOs from H9 ESCs. (**a**) A schematic depicting the main steps for hCO production. Representative bright field images of morphological changes are shown below. Triangles (Day 9) mark the inner and outer edge of the neuroepithelial ring and arrows (Day 13) indicate early ventricle structures. Scale bars: 250 μm for Day 0, 5, 9 and 13 and 1 mm for Day 60. (**b-d**) The effect of well shape and surface coating on EB formation was assessed on Day 5. (**b**) Representative bright field images of EBs generated using the indicated plate format. Scale bar = 250 μm. Non-Treated (NT), Non-Binding (NB). **(c)** Percent of cell aggregates displaying uniform density as assessed using phase-contrast microscopy is plotted as the mean ± SD (n = 3). (**d**) Individual EB diameters (black circles) and the mean (horizontal dash) ± SD (n ≥ 30/condition) is plotted. (**e**) Percent of total EBs displaying radialization neuroepithelium on Day 5 at the indicated bFGF concentrations are plotted as mean ± SD (n = 3). (**f and g**) Analysis of ventricle formation on Day 13. (**f**) Representative bright field images of COs prior to Matrigel extraction from 3 independent batches is shown. Scale bar = 500 μm. (**g**) Quantification of the diameter of individual ventricle-like structures (black circles) from three independent batches is plotted with the mean diameter marked (horizontal dash). Colored numbering corresponds to images in panel **f**. (**h**) Macroscopic organization in H9-derived hCOs. Representative bright field (Scale bar = 1 mm) images of hCOs in suspension culture at 4, 8, 12, 18, and 24 weeks of culture (top) with corresponding sections stained with DAPI to mark nuclei (bottom). Scale bar = 500 μm.

Basic FGF (bFGF) in the presence of Nodal/Activin SMAD signalling is important for maintenance of pluripotency in hES cells (Vallier, Alexander et al., 2005), whereas in the absence of SMAD signalling FGF drives neuroectoderm induction (LaVaute, Yoo et al., 2009). However, hESC aggregates display intrinsic suppression of BMP-SMAD signalling, so the addition of FGF is sufficient to drive neural induction (LaVaute et al., 2009). Furthermore, while ROCK inhibitor (Y27) promotes survival and pluripotency of singularized hESC, it is not required for EB formation (Pettinato, Wen et al., 2014). Therefore, we removed Y27 after EBs formed (2 days), and titrated FGF doses for the following 3 days to determine the optimal concentration that supports the subsequent formation of radicalized neuroepithelium following neural induction. We observed that after 3 days at 2 ng/mL, >90% of EBs displayed correctly radialized neuroepithelium (ie, a uniform, smooth clearing at the periphery: **Suppl. Fig. 1a)**, which decreased to 60% at bFGF concentrations of >6 ng/mL. In contrast, removal of bFGF strongly suppressed radialization to 20% (**Fig. 1e**), consistent with the important role of FGF signalling in neurectoderm induction observed in vivo, and in hESC aggregates (LaVaute et al., 2009, Vallier et al., 2005). hCOs develop as self-organizing systems, and EB size has a significant impact on differentiation trajectories, with smaller EBs favouring ectoderm (Bauwens, Peerani et al., 2008). Therefore, we also examined the effect of cell seeding density (1,000 to 16,000 cells/EB) on neural induction efficiency, and observed peak efficiency at 12,000 cells/well (**Suppl. Fig. 1b**). Finally, we noted that when early hCOs were extracted from Matrigel and transferred to free-floating spinning cultures, excessive Matrigel impeded the growth of morphologically uniformly-shaped hCOs. This prompted us to ensure complete removal of all Matrigel from hCOs prior to transfer to spinning culture. For this reason, we transitioned from using the “dimpled parafilm” method (Lancaster & Knoblich, 2014a) to embedding neuralized EBs in a Matrigel droplet immobilized on the surface of a 4 well plate to facilitate complete extraction. Using all of the above optimized parameters, we assessed the morphology of 18 organoids, 6 from each of three separate batches on Day 13, just prior to transfer to spinning culture. These early organoids displayed 6-10 outer ventricle-like ring structures with an average diameter of approximately 250 μm in all batches (**Fig. 1f and g**). We also noted a marked absence of the fluid filled cyst structures (**Suppl. Fig. 1c**) commonly seen with other protocols (Lancaster & Knoblich, 2014a, Lancaster et al., 2013). Thus, by optimizing early EB formation and the neural induction phases of organoid production, we reproducibly generated morphologically uniform hCOs with a cumulative efficiency of 80-90%, while restricting unwanted non-neural differentiation (**Suppl. Fig. 1d**). Similar results were also obtained using H1 ESCs (**Suppl. Fig. 1e-g**).

### Characterization of cerebral organoid development and maturation

On day 14, early organoids coated in Matrigel had a diameter of approximately 500 μm and after careful excision from the Matrigel, were transferred to spinning culture where they were left to mature as free-floating structures. By 12 weeks, hCO diameters increased to roughly 3 mm after which little additional growth was observed for up to 24 weeks of culture (**Fig. 1h**). With the exception of an occasional loss due to the fusion of 2 organoids, virtually all hCOs continued to maturity. To assess morphogenesis, we next evaluated cell type-specific marker gene expression by immunofluorescence microscopy. Distinct ventricle-like structures (ventricular units) were abundant in early stage hCOs (4-12 weeks) and diminished in later stages (18 and 24 weeks). Serial sectioning of ventricular units showed them to be spherical with a hollow center reminiscent of the apical space in ventricles in the developing cortex (**Suppl. Fig. 2a**). Staining with SOX2, which marks Radial Glial (RG) cells, showed expression in cells lining the ventricular space similar to in vivo (**Fig. 2a and Suppl. Fig. 2b**), with most ventricular units displaying robust SOX2+ staining at 4 weeks. However, by 12 weeks SOX2+ ventricular units were more variable and in older hCOs were localized to the outer regions. In contrast, NeuN, which marks mature neurons, was low in 4 weeks old hCOs, but was readily detected in hCOs 8 weeks or older, where it was present outside of the SOX2 expression domain (**Fig. 2a and Suppl. Fig. 2b**). Similar to the in vivo orientation, TUJ1, a neurofilament protein, marked RG processes running perpendicular to the ventricular zone as early as 4 weeks (**Suppl. Fig. 3a)**. Outside of the RG cells, parallel TUJ1+ processes were observed in superficially-localized neurons. The cortical layer markers, CTIP2 (gene name, *BCL11B*) and SATB2, which identify layers V (deep) and II-IV (upper), respectively, were also detected superficial to SOX2 staining in early hCOs, with CTIP2 detected as early as 4 weeks, and SATB2 first evident at 12 weeks (**Fig. 2b and Suppl. Fig. 3b**). This temporal sequence is consistent with the timing of in vivo development in which formation of deep layers precedes that of upper layer neurons. These studies further revealed that in older hCOs, while markers of RG cells and mature neurons were maintained, the cells expressing the markers became intermingled, and the organized layering of ventricular units evident in earlier hCOs was lost, as reported previously (Watanabe, Buth et al., 2017). Altogether, this analysis demonstrates that the hCOs generated using the optimized pipeline display organization and patterning reminiscent of that observed in vivo (Molnar, Clowry et al., 2019).

**Figure 2.**
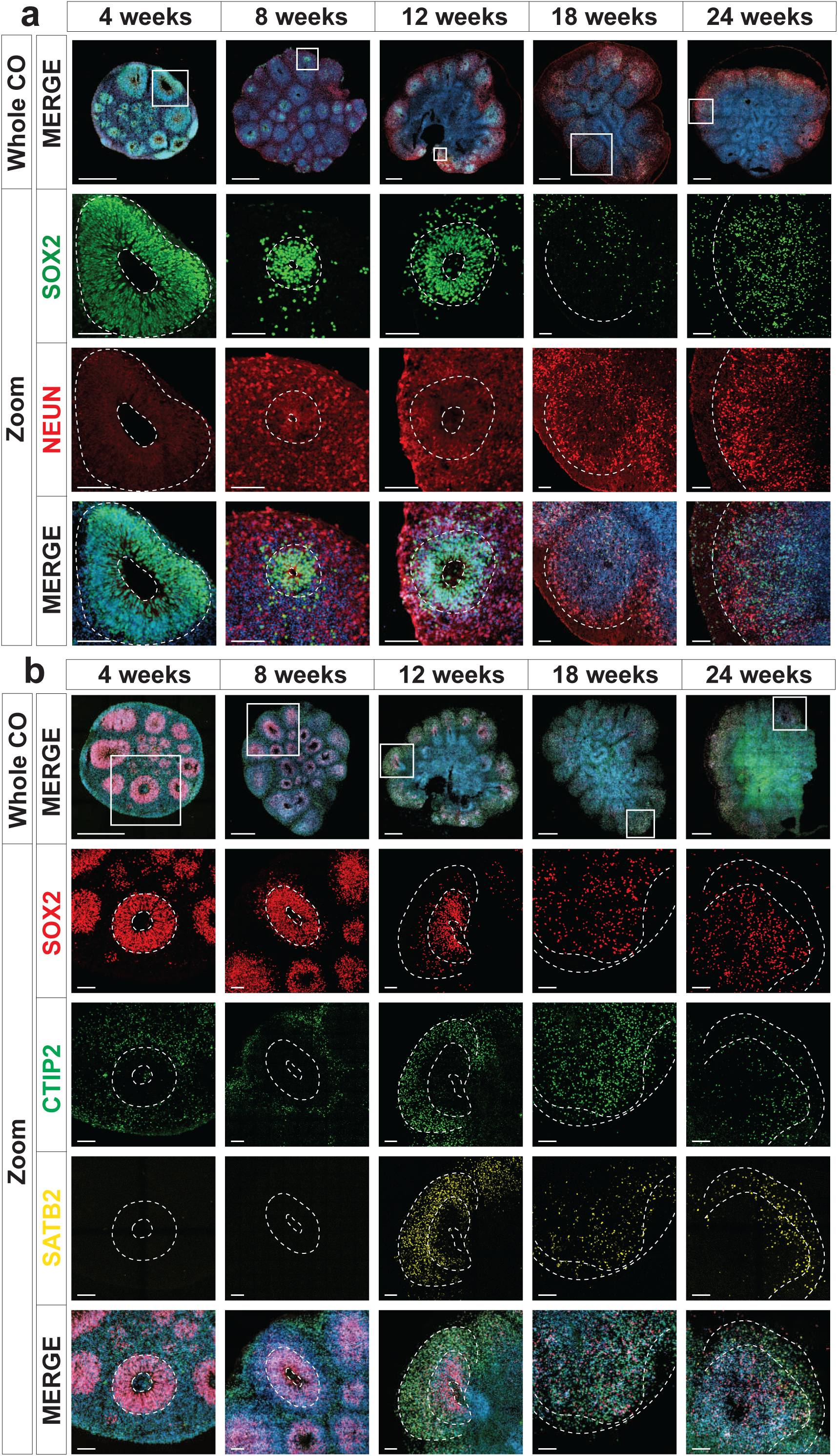
Human cerebral organoids mimics early human cortical development. The localization of SOX2 (radial glia), NeuN (neurons), CTIP2 and SATB2 (cortical layer markers) in hCOs at 4, 8, 12, 18 and 24 weeks of age, costained with DAPI was visualized by immunofluorescence microscopy. Images of whole hCOs (top) and magnified images (Zoom) of selected ventricles or regions (white box) at each time point are shown. White dashed lines mark ventricles (centre ring) and the outer perimeter SOX+ radial glial cells (outer ring). White dashed lines mark the ventricle-like cavities (inner ring) and the outer perimeter of the SOX2+ layer in 4-12 week COs or the SOX+ and outer edge of the cortical plate in 18 and 24 week COs. (**a**) Ventricle-like structures are lost in older (18 and 24 week) COs. (**b**) Expression of CTIP2 (gene name, *BCL11B*; deep layer cortical neuron marker) precedes that of SATB2 (upper layer cortical neuron marker), both of which are superficial to the SOX2+ ventricular zone in 4-12 week COs, recapitulating *in vivo* cortical development, while in older COs (18 and 24 weeks) this distinct separation is less evident. Scale bar = 500 μm for whole COs and 100 μm for magnified ventricles.

### Single Cell Profiling of hCOs

To gain a comprehensive view of cell types present in individual hCOs we performed single cell RNA sequencing (scRNA-seq) on individual organoids at 12, 18, and 24 weeks of culture. Unsupervised clustering was performed on the gene expression profiles and data was visualized using Uniform Manifold Approximation and Projection (UMAP) plots, with single cell metrics and sequencing data quality summarized in **Suppl. Table 1**. Cell types were then identified by comparing the differentially expressed genes in each of the clusters to known cell-type specific marker genes. Profiling of six organoids derived from three separate batches at 12 weeks of age identified 15 distinct clusters (**Fig. 3a and 3b and Supp. Fig. 4**). This included RG cells (*SOX2*, *PAX6*, *HES1*, and *GLI3*) of both proliferative (proRG: cluster 13, *MKI67*, *CENPF*, *TOP2A*) and outer RG (oRG; cluster 9, *HOPX*, *FAM107A*, *TNC*, *LIFR*) subtypes (Pollen, Nowakowski et al., 2015), with some overlap between these classes, consistent with RG cell function. RG cells give rise to neurons by transitioning through a transcriptionally distinct intermediate progenitor cell (IPC) state marked by the expression of *EOMES* and *PPP1R17* (Pollen et al., 2015), which mapped to cluster 1. Neurons comprised the majority (74%) of the cells, encompassing clusters 2-8, 10, and 12. These included immature neurons expressing high levels of *DCX* and *GAP43*, as well as more mature neurons that expressed deep and superficial layer markers such as *SATB2* and *BCL11B* (CTIP2). Two of the neuronal clusters (3 and 7) displayed a particularly high level of glycolytic (Gly) marker genes (*ALDOA*, *PGK1*, and *ENO1*), a neuronal subset also recently reported in hCOs (Pollen, Bhaduri et al., 2019). Interestingly, cells expressing markers of mature astrocytes (*GFAP*, *S100B*) were rare **(Fig. 3d)** suggesting neurogenesis is the primary process occurring in the first 12 weeks of culture. In addition to RG cells and neurons, we identified cluster 11 as Choroid Plexus (CP), with cells expressing characteristic CP markers (*TTR*, *AQP1*, *OTX2*, and *RSPO2*) while in one small cluster (#14) the differentially expressed genes were almost exclusively ribosomal proteins. Of note, the hCOs were comprised predominantly of dorsal forebrain neurons, as markers of midbrain and hindbrain (*HOXA3*, *HOXB3*, *IRX2*, *EN2*, *PAX2*, and *GBX2*) were not detected. Moreover, there was a complete absence of markers identifying mesenchymal lineages (*MYOG*, *MYH1*, *DCN*, and *BGN*) or retinal cells (*OPN1SW*, *RCVRN*, *TULP1*, and *ROM1*) indicating the fidelity of our optimized pipeline in promoting forebrain-specific differentiation.

**Figure 3.**
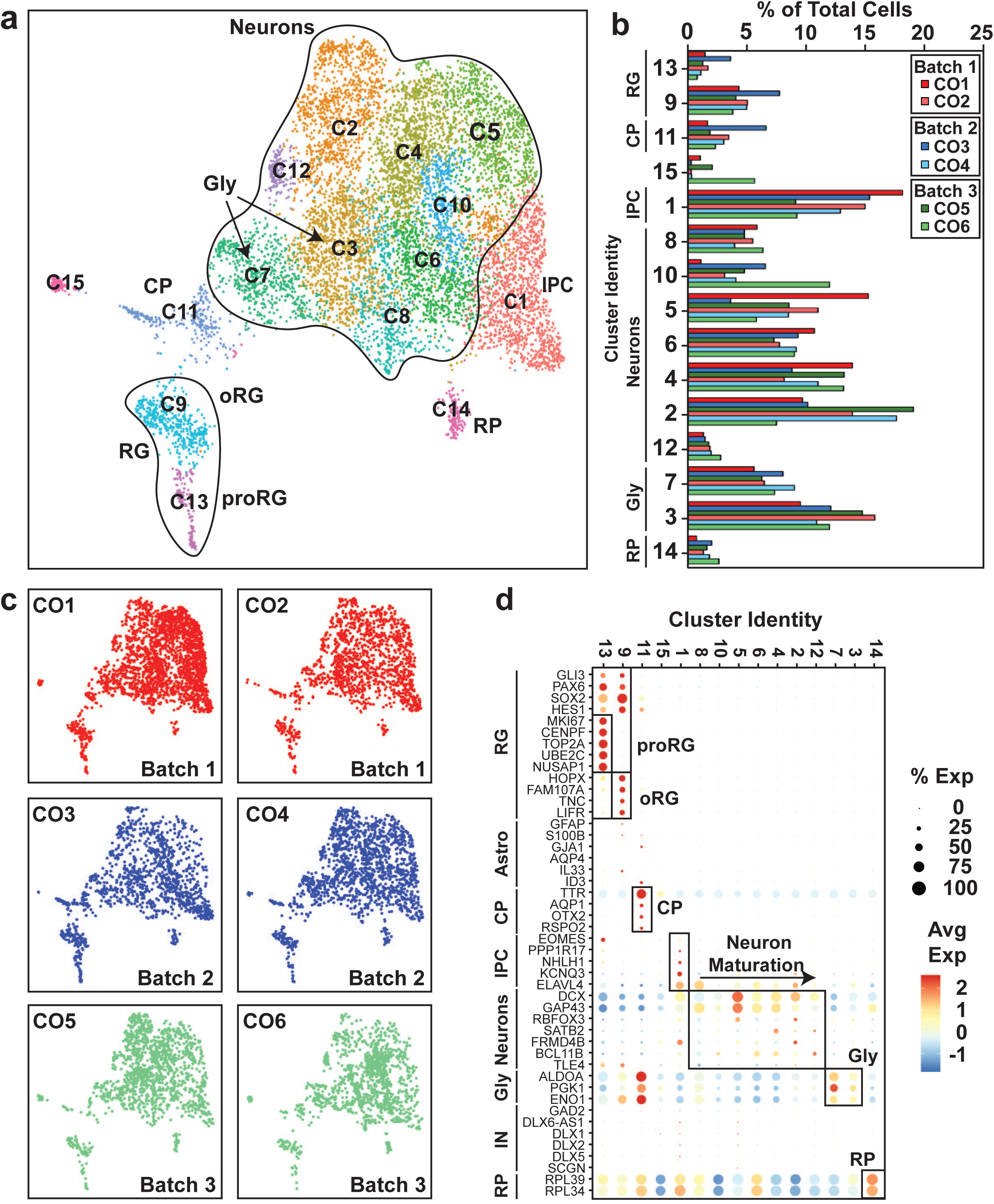
Uniform cell-type composition in 12 hCOs is revealed by scRNAseq. **(a)** A UMAP plot from unsupervised clustering of scRNAseq data from six 12 week old hCOs obtained from three separate batches (CO1-CO6) (10,985 cells) is shown. Cluster identities are indicated. **(b)** Cluster frequency analysis depicting the percentage of cells in each individual organoid that contributed to each cluster. **(c)** Individual 12 week old hCOs (CO1-CO6) are plotted in the UMAP axis defined in panel **a**. (**d**) A dot blot indicates the expression of cell-type specific marker genes for all clusters in panel **a.**

We next compared cell type composition in individual organoids to assess inter-organoid variability. For this, the percent of cells in each organoid that contributed to each of the 15 clusters was determined. This analysis revealed a remarkable conservation of cell type composition amongst the individual organoids (cluster frequency variation of <5%) even when derived from independent batches (**Fig. 3b** and **c**). Importantly, all COs contained cells from all of the clusters, indicative of conserved developmental trajectories. Transcriptional profiling of older hCOs of 18 weeks (3 hCOs from 3 batches) and 24 weeks (8 hCOs from 3 batches) was also performed (**Suppl. Fig. 5-8**). Like 12 weeks old hCOs, cell type composition was similar between individual hCOs across multiple organoids and batches at both 18 and 24 weeks, with a median cluster frequency variation of 3% and 15%, respectively (**Suppl. Fig. 5c and 7c**). Thus, even upon prolonged culturing, our optimized pipeline yields organoids of similar composition and morphology.

### Analysis of cell type diversity in maturing hCOs

To track cell composition in maturing organoids, the transcriptional profiles of 12, 18, and 24 weeks old organoids were combined and plotted in a single UMAP comprising 19 clusters (**Fig. 4a, Suppl. Fig. 9a and b and 10**). Analysis of the relative contribution of cells from each time point (**Fig. 4b, c and Suppl. Fig. 9a**) revealed the emergence at 24 weeks of a new cluster comprised of interneurons (IN; expressing *DLX1*, *DLX2*, *DLX6*, *DLX6-AS*, *GAD1*, and *GAD2*). Of note, these interneurons lacked expression of *ISL1* and *EBF1* suggesting caudal/medial identity, rather than lateral ganglionic eminence (Wamsley & Fishell, 2017). These INs cluster close to the IPC population in the UMAP, suggesting they may have arisen from these progenitors (**Fig. 3d**). We also noted another emerging subpopulation of cells within the RG/astroglia cluster (cluster 6; circle) in 24 week old hCOs (**Fig. 4a and b**). This population was marked by *OLIG1* and *OLIG2* expression, suggesting an oligodendrocyte precursor population (**Fig. 4a-d and Suppl. Fig. 9b**). RG cells are known to produce both neurons and glia, though few cells (<1%) in the 12 weeks old hCOs expressed markers of mature astrocytes (*GFAP*, *S100B*; **Fig. 4d and Suppl. Fig. 9b and c**). However, cells with an astroglial identity (Astro) were detected within the oRG cluster starting at 18 weeks (**Suppl. Fig. 5**, (cluster 5) that further increased by 24 weeks (**Suppl. Fig. 7**, cluster 3) indicating the emergence of mature astroglia from RG cells (**Fig. 4d and Suppl. Fig. 9b and c**). Overall, this suggests that the rate of gliogenesis increases upon longer culturing, which parallels the temporal regulation of cortical development in vivo (Pinto & Gotz, 2007, Sauvageot & Stiles, 2002).

**Figure 4.**
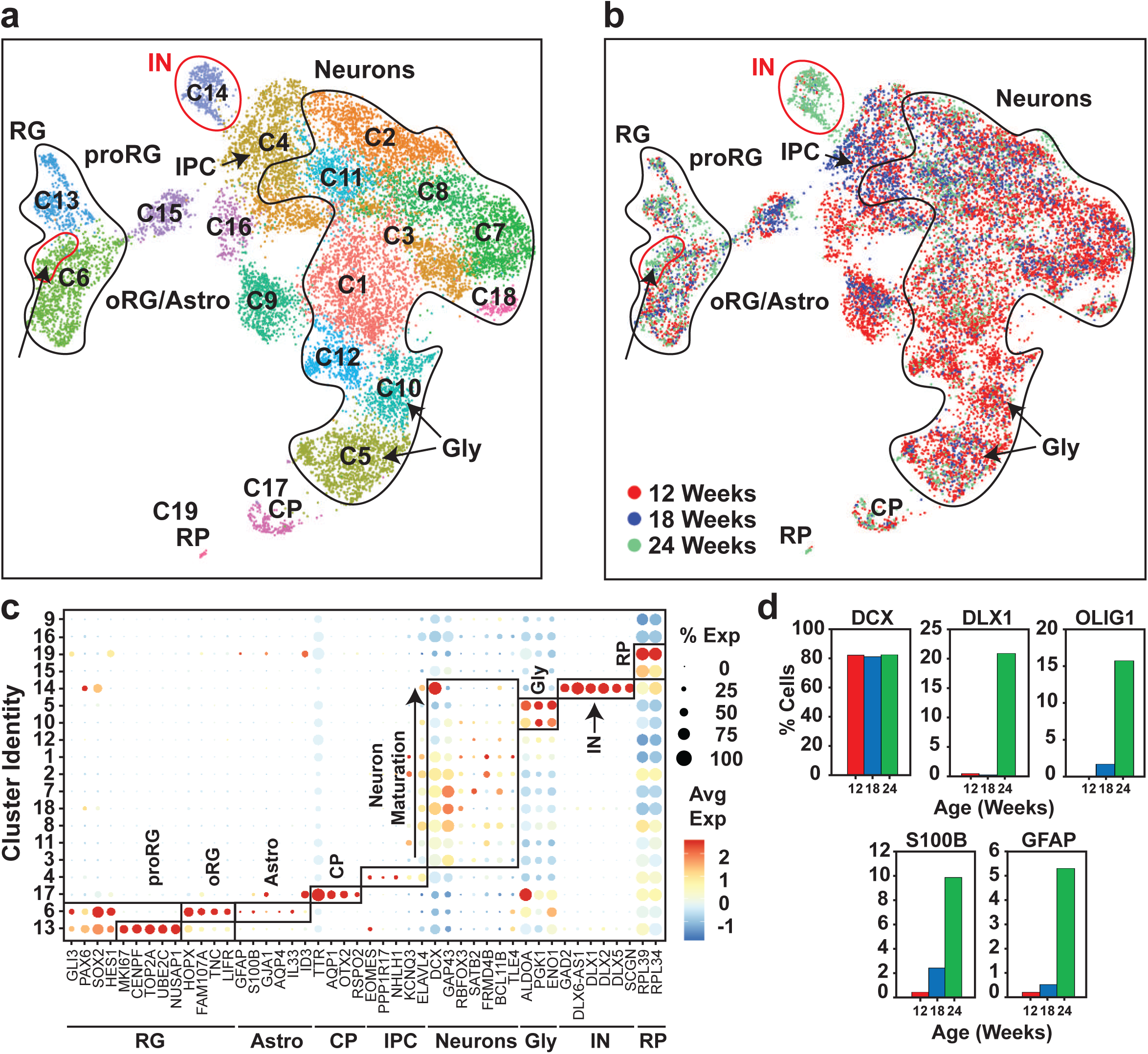
Characterization of cell type maturation in hCOs using scRNAseq. (**a**) Unsupervised clustering of combined scRNAseq data from 12, 18 and 24 week COs was visualized in UMAP plots. Cell types appearing at 24 weeks, including interneurons (IN) in cluster 14 and oligodendrocyte progenitors in oRG/Astroglial cluster 6, are circled (red). Cluster identities are indicated. (**b**) UMAP present in **a** segregated by timepoint. Cells from 12 weeks depicted in red, blue for 18 weeks and green for 24 weeks (**c**) A dot blot indicates the expression of cell-type specific marker genes for all clusters in panel **a**. The percent of cells expressing the gene (circle diameter) and the scaled average expression of the gene is indicated by the colour. (**d**) The percent of cells with expression of the indicated cell lineage markers including *DCX* (neurons), *S100B* (mature astrocytes), *GFAP* (astroglia/astrocytes), *DLX1* (interneurons) and *OLIG1* (oligodendrocyte precursors) and at 12, 18 and 24 week time points is plotted. Note the general neuronal marker *DCX*, which is expressed similarly across the time points was used as a reference. RG, radial glial cells; oRG, outer radial glial cells; proRG, proliferative radial glial; CP, choroid plexus; IPC, intermediate progenitor cells; GLY, glycolytic signature; RP, Ribosomal Protein; IN, interneurons.

### Electrophysiological Analysis

To assess neuronal function, electrophysiological output was measured by whole cell patch clamping of individual neurons in fresh slices prepared from hCOs at 12 and 24 weeks. Patched neurons displayed varying degrees of neurite networks as visualized by biocytin labelling (**Fig. 5a**). Analysis of electrical properties identified 3 types of neurons, namely immature, developing, and mature. Immature neurons did not fire action potentials (AP; data not shown), developing neurons, fired APs but with slow kinetics and small amplitudes, and mature neurons, fired APs with fast kinetics and high amplitudes and generated stable trains of spontaneous APs (**Fig. 5b and c**). Both developing and mature neurons also displayed sodium and potassium currents though they were smaller in the developing neurons (**Suppl. Fig. 11**). Moreover, only mature neurons generated stable spontaneous APs upon slight depolarization though both developing and mature neurons displayed similar spontaneous AP frequency and amplitude (**Fig. 5d-f**). Of note, all three neuron classes were present in both 12 and 24 weeks old hCOs, indicating that even at 12 weeks, electrophysiologically mature neurons were present.

**Figure 5.**
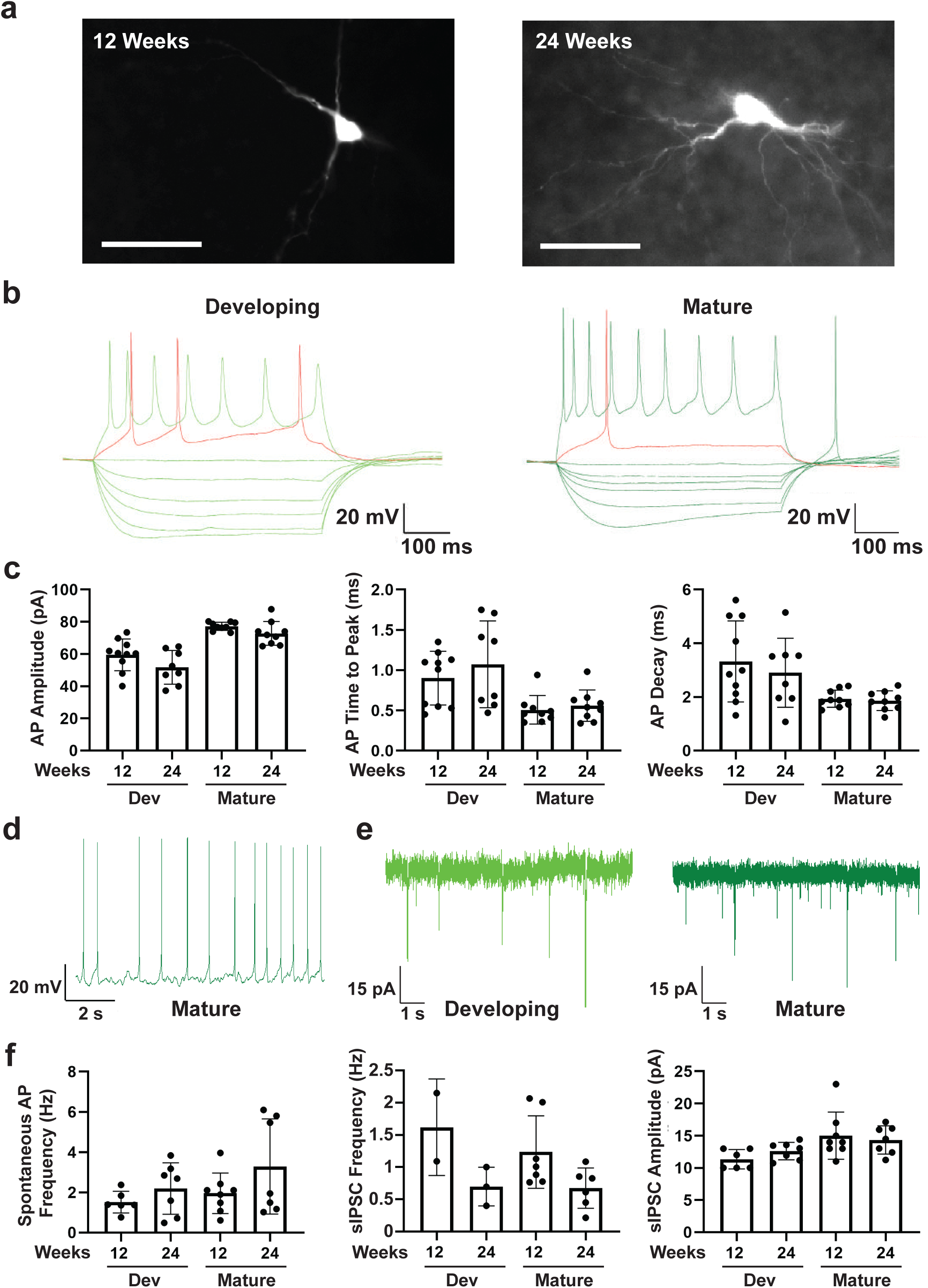
Electrophysiological analysis of 12 and 24 week old hCOs using whole cell patch clamping. (**a**) Images of recorded developing or mature neurons marked with biocytin taken using 40x water immersion objective are shown. Scale bar = 20 μm. (**b**) Representative traces from whole cell patch clamping of developing and mature neurons are shown. **(c)** Peak action potential amplitude ± SD, mean time to peak action potential amplitude ± SEM, and mean action potential decay time ± SEM, for individual recordings are plotted for 8-10 neurons per condition. **(d-e)** Characterization of spontaneous currents. A representative trace of spontaneous AP firing in a mature neuron **(d)** and of mean frequency ± SEM of spontaneous excitatory postsynaptic currents (EPSCs) in developing and mature neurons (**e**) are shown. (**f**) Frequency and amplitude of spontaneous IPSCs in developing and mature neurons is plotted as the mean ± SEM for 2-6 neurons per condition.

## Discussion

Human stem cell derived cerebral organoids provide an unparalleled model system to study human neocortical development and associated disease processes. However, a key limitation for research applications is the considerable variability in shape/architecture and cell type composition present in individual organoids. This characteristic makes it particularly challenging to design experiments to address the effects of genetic variants, therapeutic candidates and other perturbations affect CO morphogenesis or function such that allow statistically supported conclusions to be drawn. Here, we describe a protocol to efficiently generate human dorsal forebrain organoids with phenotypically uniform morphologies that are reproducibly comprised of similar proportions of cell types across independent batches. This highly reproducible model system is thus amenable for the study of pathways controlling early human brain development and disease processes.

Several protocols have been developed to generate cerebral organoids including both directed and self-directed (Kadoshima, Sakaguchi et al., 2013, Velasco et al., 2019, Yoon et al., 2019) and here we selected a previously established method of self-patterned organoids (Lancaster & Knoblich, 2014a, Lancaster et al., 2013) as a starting point. Our work to standardize production revealed that optimization of the early steps of organoid generation including the embryoid body formation and neural induction phases were key in ensuring the production of similar COs. This included normalizing EB morphogenesis using early removal of ROCKi and optimized FGF treatment, cell numbers, and well geometry and coatings. Finally, we found that efficient removal of Matrigel prior to spinning culture improved subsequent development. Applying this method, we demonstrated using single cell profiling that our individual COs have extensive cell diversity but with similar proportions of cell types even across different batches. In contrast, the reported single cell analysis of the originally established method revealed considerable variability in the proportions of cell types present in individual organoids, particularly across different batches (Quadrato et al., 2017). Furthermore, we observed uniform distribution of ventricular units, and that the cell type composition evident in 12 week old hCOs was maintained in older organoids, with additional cell types, including interneurons, mature astrocytes and oligodendrocyte precursors emerging at 24 weeks. Importantly, these new cell types were present in all organoids at similar proportions. Finally, our electrophysiological analysis revealed the presence of mature, electrically active neurons.

Our optimized self-directed protocol that generates highly reproducible hCOs, complements those reported in other recent studies that use a directed patterning approach involving small molecule inhibitors and growth factors (Kadoshima, Sakaguchi et al., 2013, Velasco et al., 2019, Yoon et al., 2019). The availability of several robust platforms to produce highly similar hCOs, including the one described herein, thus allows for systematic molecular and cellular characterization of the development of the human cortex and how disease-causing mutations alter development and perhaps also homeostatic events.

## Materials and Methods

### Cell line generation and culturing

H9 and H1 ESCs were cultured on plates coated with hESC grade Matrigel (VWR, #CA89050-192) at 37°C and 5% CO_2_ in mitotically arrested Mouse Embryonic Fibroblasts (MEFs) conditioned media (DMEM/F12 (Life Technologies, #11330057), 20% Knock-out Serum Replacement (KSR: Life Technologies, #A3181502-02), 2% MEM-NEAA, 55 μM β-Mercaptoethanol (Life Technologies, #21985-023), and 4 ng/mL basic fibroblast growth factor (bFGF; Peprotech, #100-18B). Media was changed daily and cells were split at a 1:6 ratio using Collagenase IV (Stemcell Technologies, #07909) every 5-6 days.

### Generation of Cerebral Organoids

ESCs were singularized using TrypLe Select (Life Technologies, #12563011), and resuspended at 80,000 cells/mL in Embryoid Body (EB) media (DMEM/F12, 20% KSR, 2% MEM-NEAA, 55 μM β-Mercaptoethanol) with 4 ng/mL bFGF and 50 μM Y-27632 (Y27) (Selleck Chem, #S1049). Cells (12,000 cells/well) were plated in 96 well “V” bottom non-binding plates (Greiner Bio-One, #651970) and on day 2, fresh EB media containing 2 ng/mL bFGF was added. On day 5, healthy EBs with a diameter of 425-475 μm were transferred to 24 well ultra-low attachment plates (Corning, #CLS3473) in 500 μL neural induction media (DMEM/F12, 1% N2, 1% MEM-NEAA, 1% Glutamax, and 1 μg/mL heparin (Sigma, #H3393)), and 48 h later an additional 500 μL of neural induction media was added. EBs were transferred to pre-warmed 4 well tissue culture plates, excess media was aspirated and freshly thawed growth factor reduced Matrigel (30 μl) was added on top of EBs. Plates were transferred to a 37°C CO_2_ incubator for 10 minutes to allow the Matrigel to polymerize and then 500 μL of Cerebral Organoid Differentiation Media without vitamin A (CDM-Vit A; 48% DMEM/F12, 48% Neurobasal (Life Technologies, #21103049), 0.5% MEM-NEAA, 1% Glutamax, 0.5% N2, 1% B27 without vitamin A (Life Technologies, #12587001), 2.5 μM insulin, and 192.5 μM β-Mercaptoethanol was added to each well. On day 11, media was replaced and on day 13, spheroids containing ring-like structures were extracted using a Scalpel (ThermoFisher, #1000044), ensuring that excess Matrigel is removed. Organoids (maximum of 3 per well) were transferred to a 6 well tissue culture plate containing 3 mL of CDM + Vit A (CDM with 1% B27 containing Vit A), and then placed on an orbital shaker (ThermoFisher, #88881101) at 90 RPM in a CO_2_ incubator for the duration of culturing. Media (2 mL) was replaced every 72 hours for the first 30 days and then increased to 3 mL for the duration of culture. Plates were replaced every 30 days to prevent build up of debris. Spheroid formation efficiency was tested using non-binding V bottom plates (Bio-One, #651970), Aggrewells coated with anti-adherence rinsing solution (StemCell Tech), U bottom non-treated polystyrene plates (ThermoFisher #168136) and U bottom, ultra low attachment plates (Corning#4515).

### Immunofluorescence Microscopy

COs were washed 3 times for 5 minutes in 5 mL of PBS in 5 mL round bottom tubes (BD Falcon, #1152367) then fixed overnight in 4% PFA at 4°C on a rocker. Samples were washed 3 times for 5 minutes in 0.1% Tween20 in PBS (PBST) and then sequentially soaked in 15% sucrose solution for 2 hours and then 30% sucrose solution overnight (4°C). Organoids were transferred to a cryo mold containing OCT (VWR, #95057-838) and flash frozen in liquid nitrogen using a stainless steel bucket containing 2-methylbutane (Sigma, #M32631). Samples were sectioned to 20 μm using a Leica CM3050S cryostat, mounted on Superfrost Plus Microscope Slides (Fisher Scientific, #22-037-246) and heated for 15 minutes at 45°C. Samples were washed 3 times in PBST a room temperature, blocked and permeabilized in for 1 hour at room temperature in 2% BSA and 0.5% Triton-X100 diluted in PBST (PBSTx) in a humidified chamber. Primary antibodies listed in Suppl. Table 2 were diluted in cold PBST containing 0.5% BSA and incubated overnight at 4°C washed 3 times for 5 minutes in PBST at room temperature and then incubated in Invitrogen DyLight™ Secondary Antibodies (1:500) and DAPI (1:2000) in PBST containing 0.5% BSA overnight at 4°C. Slides were washed in PBST 3 times for 10 minutes and mounted using MOWIOL-DABCO Mounting Media (Sigma, #10891) and then dried at room temperature overnight in a dark box. Samples were imaged using a Nikon T2i microscope with a Hamamatsu confocal camera or Zeiss CSU-X1 confocal microscope. Images were processed using NIS elements (Nikon Elements) software or Volocity (PerkinElmer) software, respectively.

### Paraffin embedding

COs were washed 2 times for 5 minutes at room temperature in 5 mL of PBS in 5 mL round bottom tubes (BD Falcon, #1152367) then fixed overnight in 4% PFA at 4°C on a rocker. Samples were washed 3 times for 15 minutes in PBS and then sequentially dehydrated in 40%, 70%, and 95% ethanol diluted in ddH_2_O for 1 hour each, 95% ethanol overnight and then 3 times for 1 h in 100% ethanol at 4°C on a rocker. COs were then soaked in Xylene for 5-10 minutes at room temperature in biopsy cassettes (Simport, #M506-3). Samples were air dried for 2-3 minutes, washed twice in melted Paraffin (Leica, #39602004) at 60 °C for 1 hour and once overnight. Samples were embedded into paraffin blocks within 16 hours and then sectioned using 80 mm microtome blades (Thermo Scientific, MB35) at 14 or 20 μm on a semi-automated rotary microtome (Leica, RM2245) and mounted on Superfrost Plus Microscope Slides in distilled H_2_O heated to 45°C, then dried over night at 45°C and stored at 4 °C.

### Hematoxylin and eosin staining

Paraffin slides were deparaffinized by 2 washes of Xylene, 100%, 100%, 95%, and 70% ethanol for 5 minutes each. Deparaffinized or cryopreserved slides were soaked for 5 min in dH_2_O, stained with Harris’ Hematoxylin (Sigma, #HHS32) for 3 minutes, rinsed in warm running tap water for 4 minutes, and then destained in 0.3% acid ethanol (70% ethanol and 0.3% HCl) for 30 seconds and rinsed in warm running tap water for 5 minutes. Samples were dipped in bluing agent 0.1% sodium carbonate (Sigma, #1613757), washed in running tap water for 2 minutes and then counterstained in a working solution of 0.25% eosin (in 80% ethanol and 0.5% glacial acetic acid) for 20 seconds. Slides were sequentially dehydrated for 1 min in 70%, 95%, and 100% ethanol and for 5 minutes in Xylene or Histo-Clear II (National Diagnositics, #HS-202), mounted using Histomount (Thermofisher, #008030) and left to dry overnight prior to imaging. H&E images were taken on a dissection scope with a mounted camera, or with a 20x objective on Axio Scan.Z1 (Zeiss) and processed using Zen (Zeiss) software.

### Sample preparation for scRNA-seq

COs were washed 3 times for 1 minute each in with 5 mL of room temperature PBS, cut into fragments, digested with Accutase at 37°C for 20 minutes, with pipetting every 10 minutes to facilitate cell dissociation. The cell suspension was passed through a 40 micron filter, which was then washed once with 2 mL of fresh Accutase to collect remaining single cells and then centrifuged at 700 RPM. The pellet was resuspended in 5 mL of Accutase at room temperature and live cells counted using trypan blue staining, centrifuged at 700 RPM for 5 min and single cells resuspended in CDM, +Vit A at 500,000-1,000,000 cells/mL and trypan blue positive cells recounted. Samples were placed on ice and then subjected to barcoding using the 10X Chromium Platform following the manufacturer’s recommendations. cDNA libraries were sequenced on either Illumina’s HiSeq 3000 or a NovaSeq 6000.

### Analysis of scRNA-seq data

Single cell RNA sequences were processed with Cellranger (V2.1) software (10X Genomics) to perform demultiplexing, UMI (unique molecular identifier) collapsing, demultiplexing, and alignment to the GRCh38 human transcriptome. Post-processing of the raw cellranger output matrix was performed using an in-house pipeline (Ayyaz, Kumar et al., 2019). Data processing including cell filtering using a stringent filter, gene filtering, and data normalization for individual samples were performed with scater (McCarthy, Campbell et al., 2017). Merging of multiple samples into a combined object, unsupervised clustering, visualizations, and differential expression analysis was done using Seurat (Butler, Hoffman et al., 2018, Stuart, Butler et al., 2019). In Seurat, UMAP was used as the preferred dimensionality reduction method (Becht, McInnes et al., 2018) whereas MAST was used for differential expression analysis (Finak, McDavid et al., 2015). Batch correction between multiple batches in the same time point as well as between time points was performed using Harmony (Korsunsky, Fan et al., 2018). All data plotting and analyses were done in R (www.R-project.org).

### Electrophysiological recordings

CO (500 μm) were embedded in 0.4% low melting agarose (Thermofisher, #16520100) diluted in PBS and then sectioned using a Leica VT1200S vibratome. Slices were perfused at a constant rate of 2 mL/min with oxygenated, warmed recording artificial cerebrospinal fluid solution, containing 124 mM NaCl, 25 mM NaHCO_3_, 2.5 mM KCl, 2.5 mM MgCl_2_, 1.2 mM CaCl_2_, and 10 mM glucose at a temperature 32 ± 1°C. Visually guided whole-cell patch-clamp recordings were obtained from neurons with a patch solution containing: 120 mM K-gluconate, 20 mM KCl, 10 mM HEPES, 2 mM MgCl_2_, 2 mM Mg_2_ATP, 0.3 mM NaGTP, 7 mM phosphocreatine, 0.6 mM EGTA (pH = 7.2, 295 mOsm). Biocytin was routinely added to patch solution to reveal morphology of recorded neurons. Immediately after recordings, slices were fixed in 4% PFA and immunostaining for biocytin was done using streptavidin-Alexa 546 conjugated antibodies as described previously (Khlghatyan, Quintana et al., 2019).

## Data availability

The accession number for the gene expression data reported in this paper is GEO: GSE137877.

## Acknowledgements

This work was supported by funds from CFREF/Medicine by Design, Krembil Foundation and Canadian Institute for Health Research (CIHR) Foundation grants to L.A. and J.L.W. and from CIHR and Canada Research Chair to J.M.B. The authors have no conflicts of interest to declare.

## Author Contributions

A.S., J.T.G., J.L.W. and L.A. conceived the project and wrote the manuscript. A.S, J.T.G., B.G., A.E., D.T., J.J.H. and P.G. performed experiments. L.A., J.L.W., A.S. and J.T.G. planned and organized the work. J.M.B., J.L.W and L.A. secured funding.

## SUPPLEMENTARY INFORMATION

**Supplementary Figure 1.**
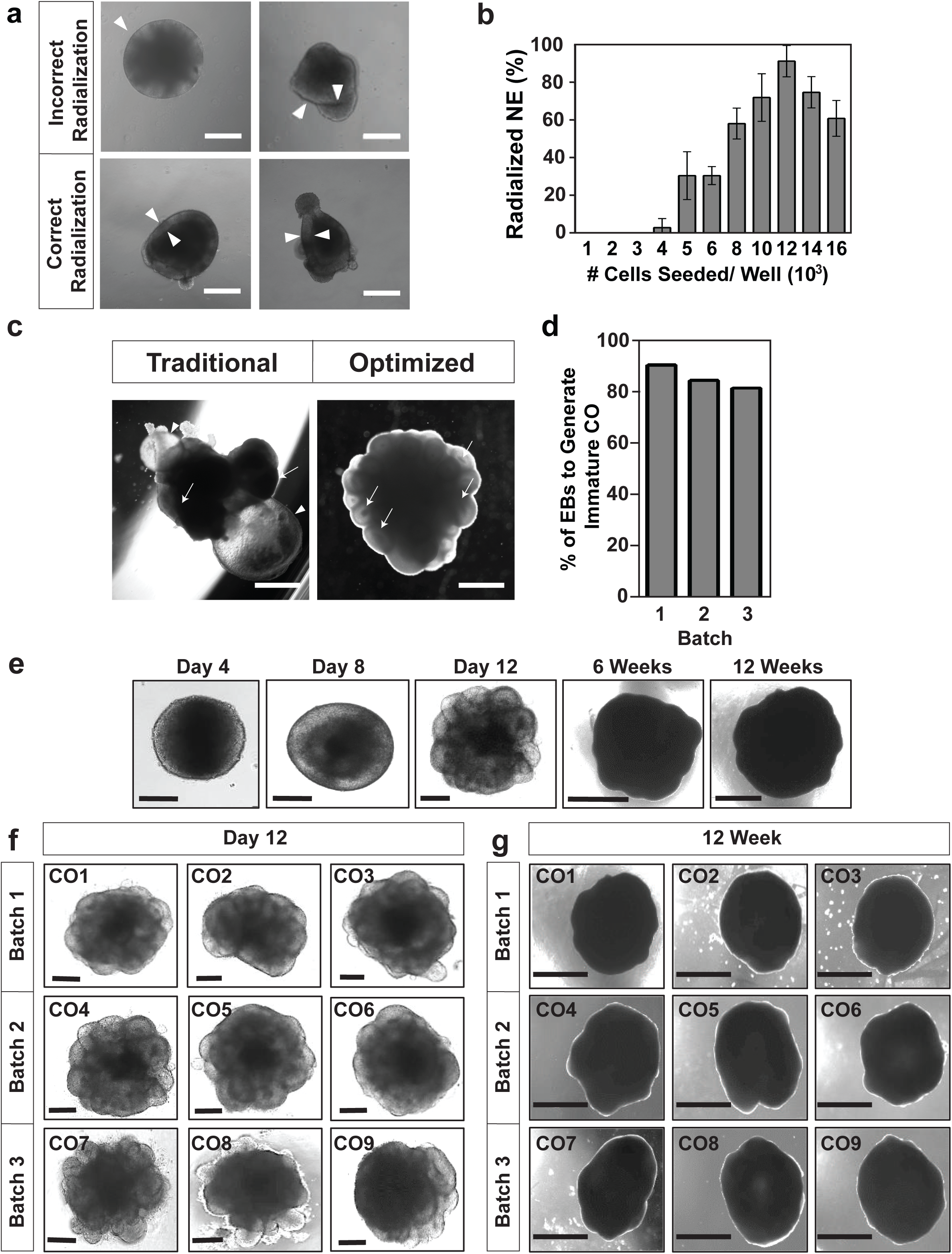
Morphological characterization of H9 and H1-derived hCOs. (**a**) Representative bright field images of failed (top panels) or successful (bottom panels) neuroepithelialization on Day 4 of neural induction from H9 ESCs. White triangles denote the neuroepithelial ring. Scale bar = 250 μm. (**b**) Percent of total EBs displaying radialization neuroepithelium on Day 5 at the indicated cell seeding concentrations are plotted as mean ± SD (n = 3). (**c**) Representative images of hCOs generated using the optimized and traditional CO differentiation pipelines. Arrows show ventricle structures and the arrowhead highlights a fluid filled cyst structure on the organoid exterior. Scale bar = 1 mm. (**d**) Efficiency of H9-derived COs displaying ventricle ring structures on Day 13, prior to Matrigel extraction, from three independent batches of COs. (**e-g**). Generation of H1-derived hCOs. (**e**) Representative bright field images of morphological changes at the indicated ages are shown. (**f,g**) Representative bright field images of COs prior to Matrigel extraction on Day 12 (**f**) and at 12 weeks (**g**) from 3 independent batches are shown. Scale bars: 200 μm for Day 4-12 and 1 mm for 8 and 12 weeks.

**Supplementary Figure 2.**
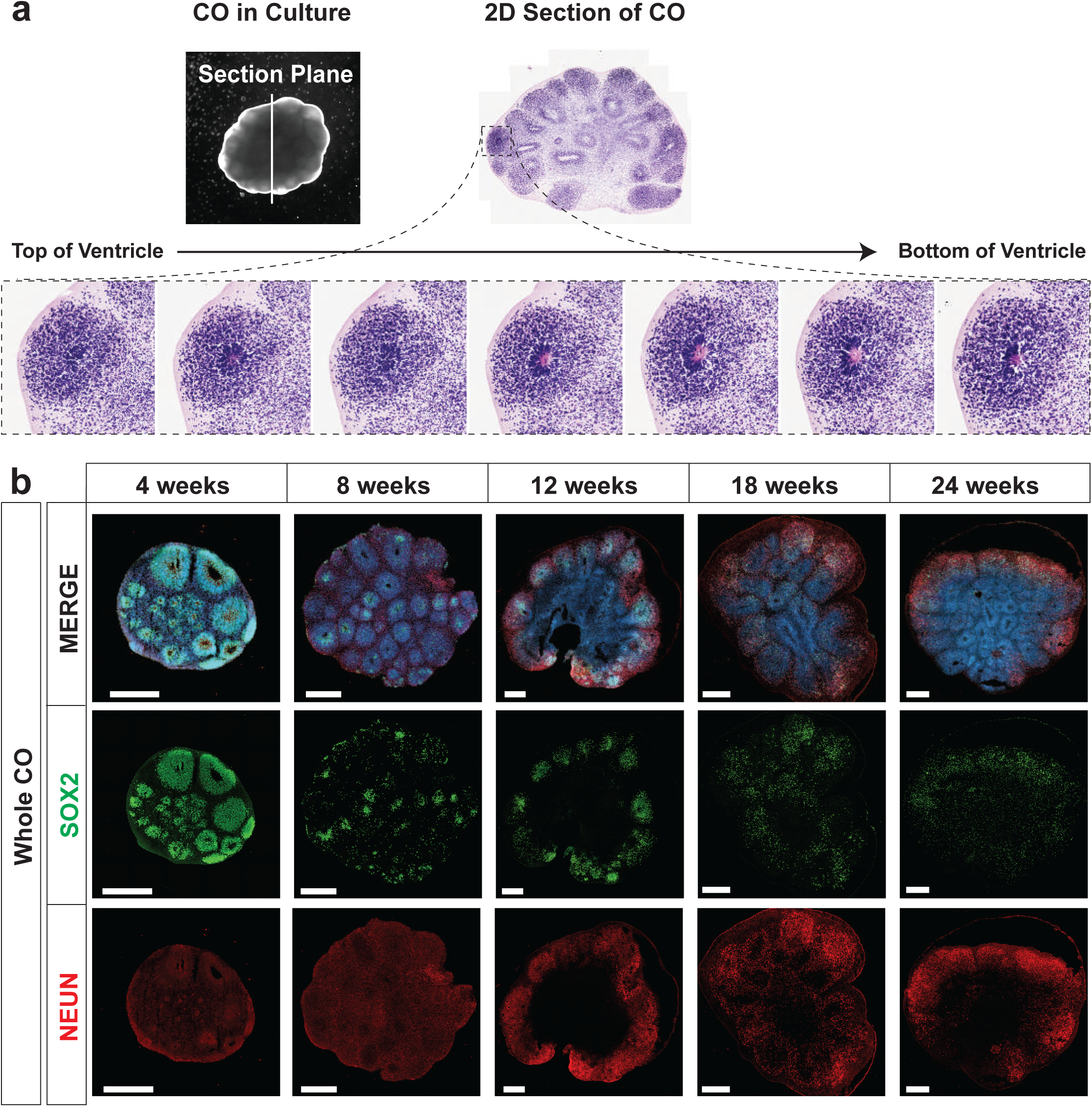
Human cerebral organoids mimics early human cortical development. (**a**) Orientation of ventricle unit in hCOs in 3D space. Visualization of serial sections of a hCO and a top view of a sectioned ventricle when mounted on a glass coverslip (top). H&E staining of 7 serial sections of a ventricle to reconstruct the 3D structure (bottom). (**b**) Images of individual channels of the merged whole CO images shown in Figure 2. The localization of SOX2 (radial glia), NeuN (neurons) in hCOs at 4, 8, 12, 18, and 24 weeks of age, costained with DAPI was visualized by immunofluorescence microscopy. Scale bar = 500 μm for Whole COs.

**Supplementary Figure 3.**
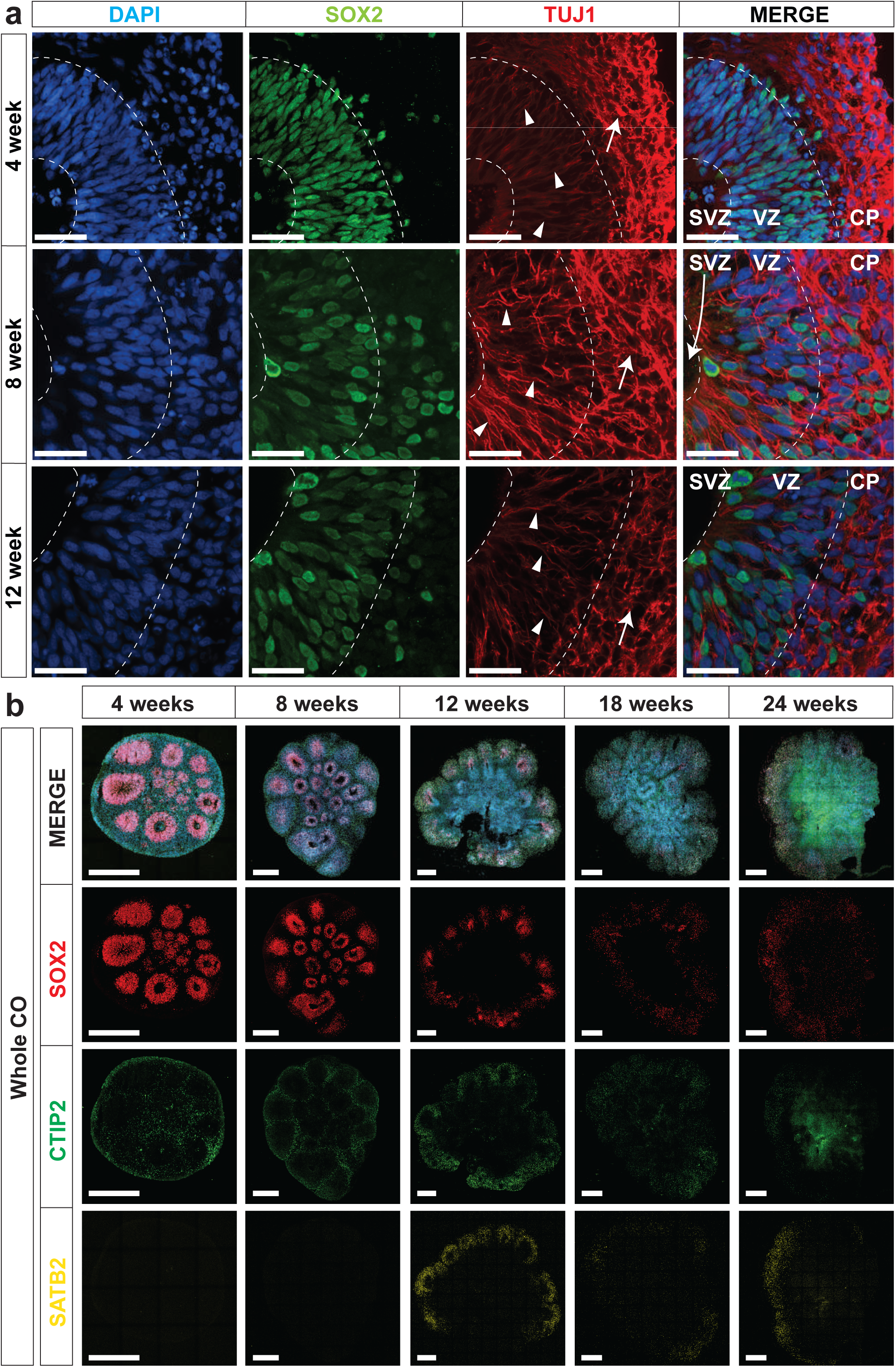
Characterization of ventricle-like structures in hCOs. (**a**) The localization of SOX2 (radial glia) and TUJ1 (neurons) in hCOs at 4, 8, and 12 weeks of age, costained with DAPI, was visualized by immunofluorescence microscopy. SOX2+ radial glial cells line the apical space of the ventricular zone adjacent to the hollow ventricle structure. Triangles denote TUJ1+ radial glial processes oriented perpendicular in the ventricular zone (VZ) and arrows denote TUJ1+ neurons with parallel processes in the cortical plate (CP). The dashed line denotes layering present in hCOs. SVZ, subventricular zone; VZ, ventricular zone; and CP, cortical plate. Scale bar = 50 μm. (**b**) The localization of SOX2 (radial glia), CTIP2, and SATB2 (cortical layer markers) in hCOs at 4, 8, 12, 18, and 24 weeks of age, costained with DAPI was visualized by immunofluorescence microscopy. Scale bar = 500 µm for Whole COs.

**Supplementary Figure 4.**
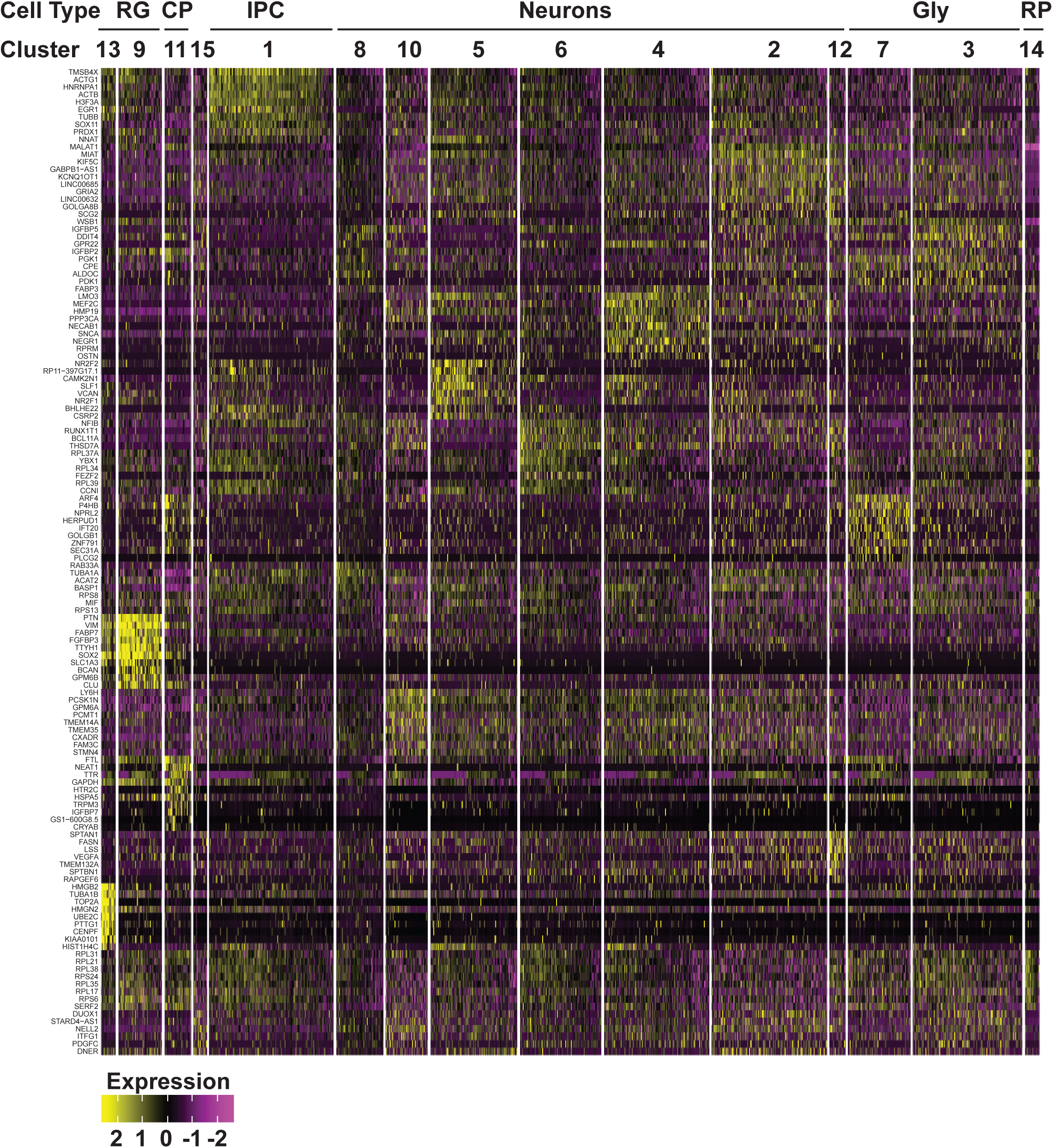
Gene expression heat map from 12 week old COs. The top 10 most differentially expressed genes per cluster are shown.

**Supplementary Figure 5.**
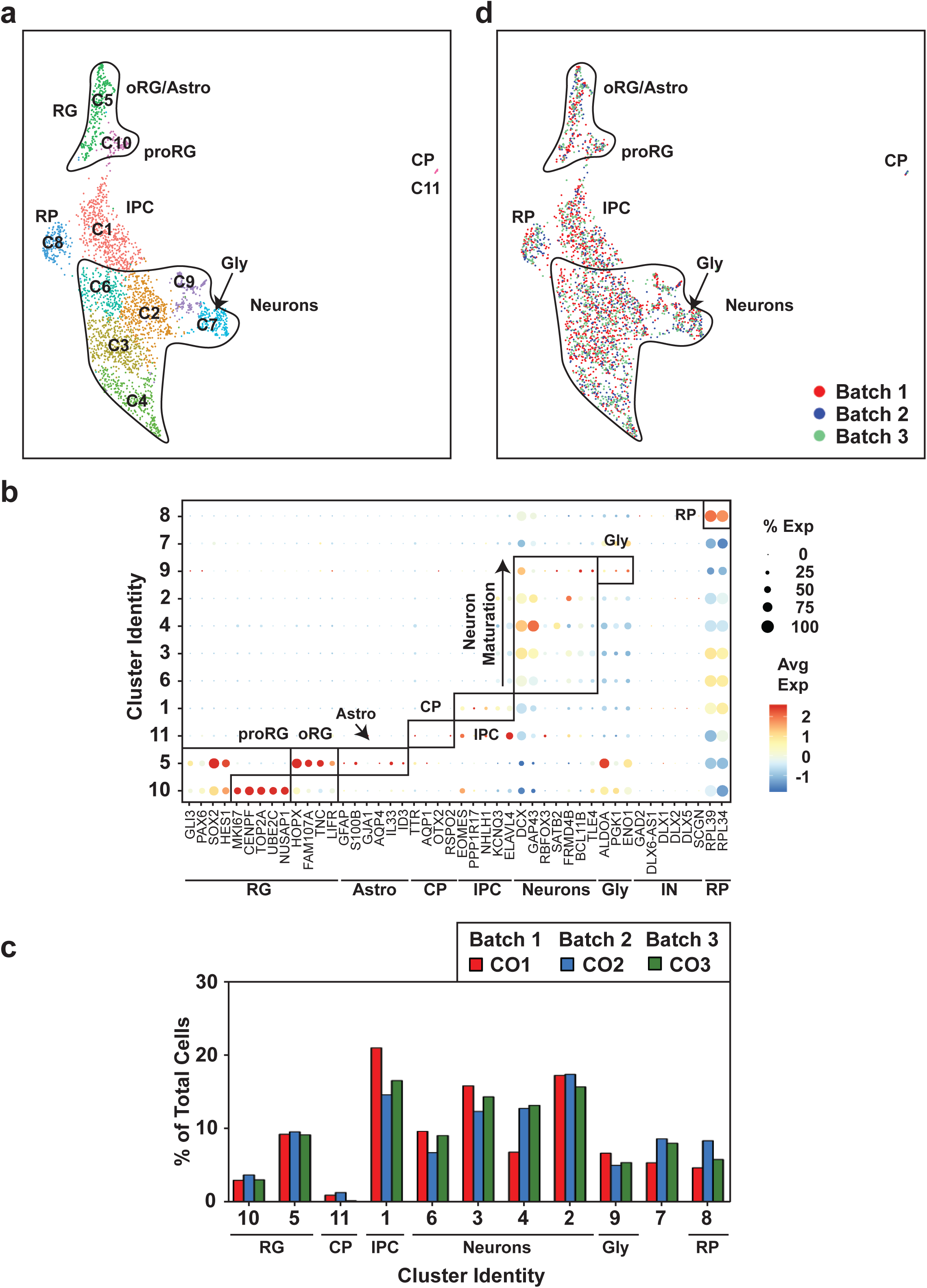
Uniform cell-type composition in 18 week hCOs is revealed by scRNAseq. **(a)** UMAP plots from unsupervised clustering of scRNAseq data from three hCOs from three separate batches (CO1-CO3) (12,000 cells) is shown. Cluster identities are indicated. The emergence of an astrocyte population is observed in oRG/astro cluster 10. **(b)** A dot blot indicates the expression of cell-type specific marker genes for all clusters. The percent of cells expressing the gene (circle diameter) and the scaled average expression of the gene is indicated by the colour. **(c)** Cluster frequency analysis depicting the percentage of cells in each individual organoid that contributed to each cluster. **(d)** Individual hCOs (CO1-CO3) are plotted in the UMAP axis defined in panel **a**. RG, radial glial cells; oRG/astro, outer radial glial cells/astroglia; proRG, proliferative radial glial; CP, choroid plexus; IPC, intermediate progenitor cells; GLY, glycolytic signature; RP, Ribosomal Protein.

**Supplementary Figure 6.**
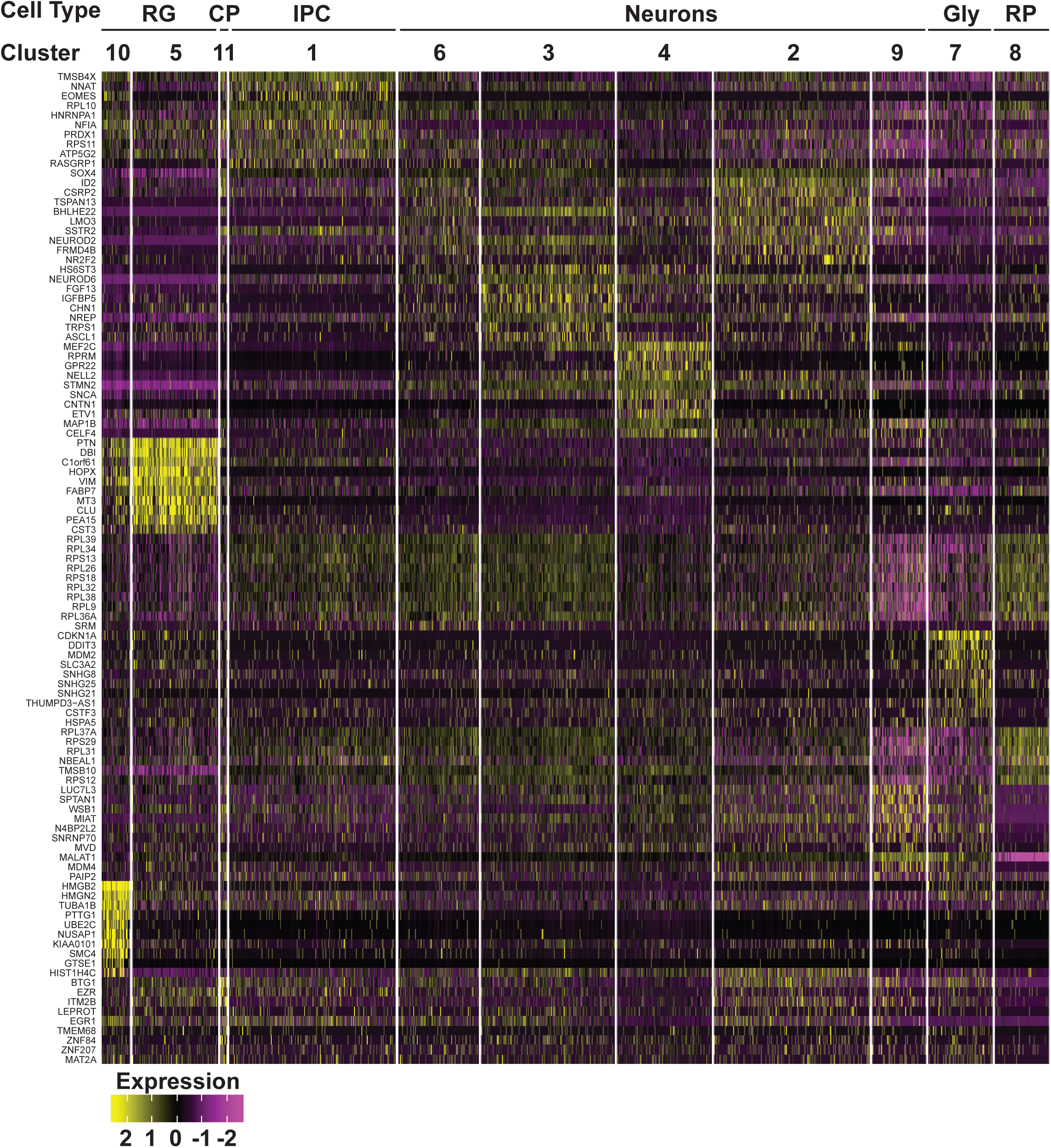
Gene expression heat map from 18 week old COs. The top 10 most differentially expressed genes per cluster are shown.

**Supplementary Figure 7.**
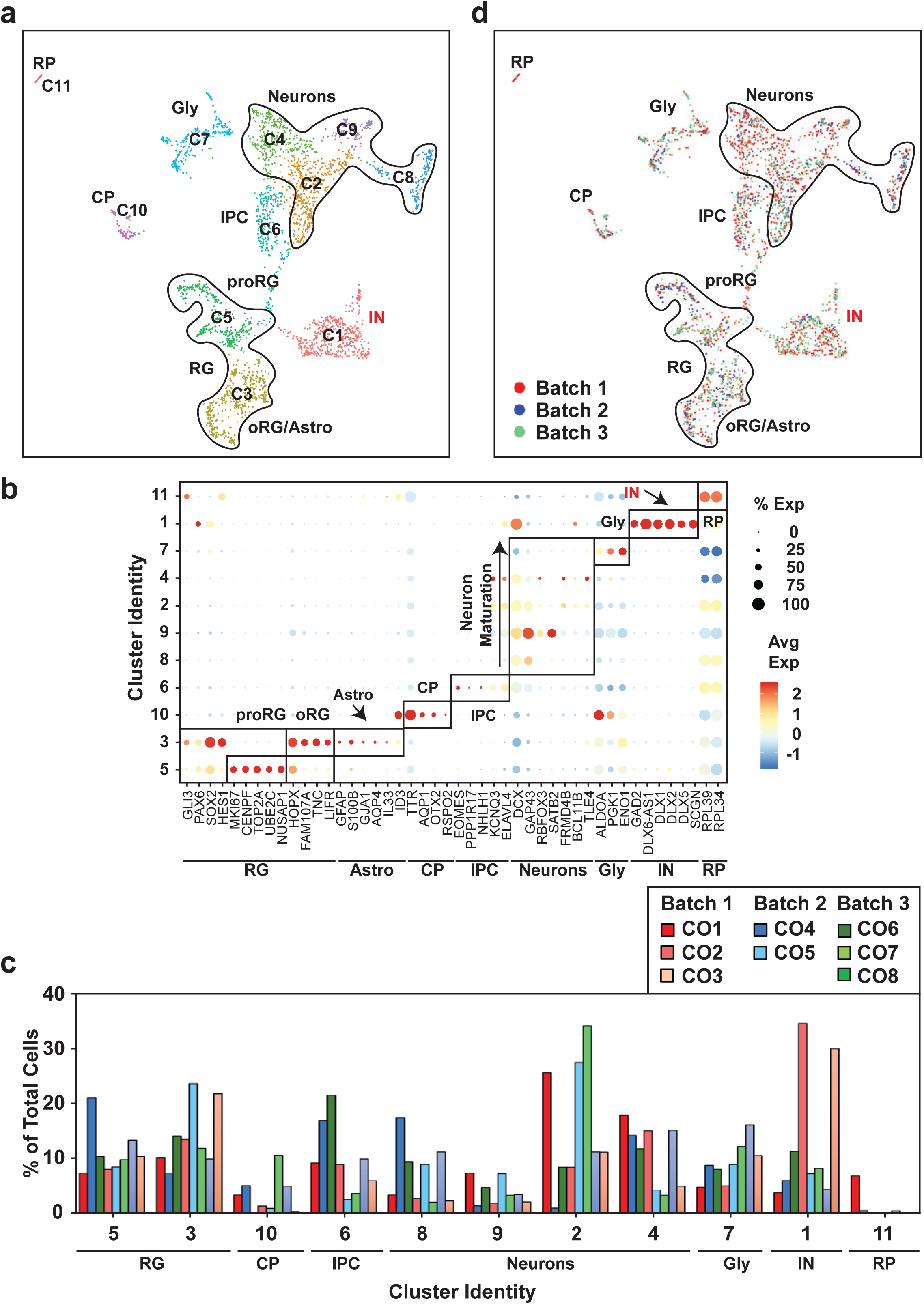
Uniform cell-type composition in 24 week hCOs is revealed by scRNAseq. **(a)** UMAP plots from unsupervised clustering of scRNAseq data from eight hCOs from three separate batches (CO1-CO8) (3,140 cells) is shown. Cluster identities are indicated. The emergence of an interneuron (IN; red text) and mature astrocyte population is observed. **(b)** A dot blot indicates the expression of cell-type specific marker genes for all clusters. The percent of cells expressing the gene (circle diameter) and the scaled average expression of the gene is indicated by the colour. **(c)** Cluster frequency analysis depicting the percentage of cells in each individual organoid that contributed to each cluster. **(d)** COs derived from each independent batch are plotted in the UMAP axis defined in panel **a**. RG, radial glial cells; oRG/astro, outer radial glial cells/astroglia; proRG, proliferative radial glial; CP, choroid plexus; IPC, intermediate progenitor cells; GLY, glycolytic signature; RP, Ribosomal Protein; IN, interneurons.

**Supplementary Figure 8.**
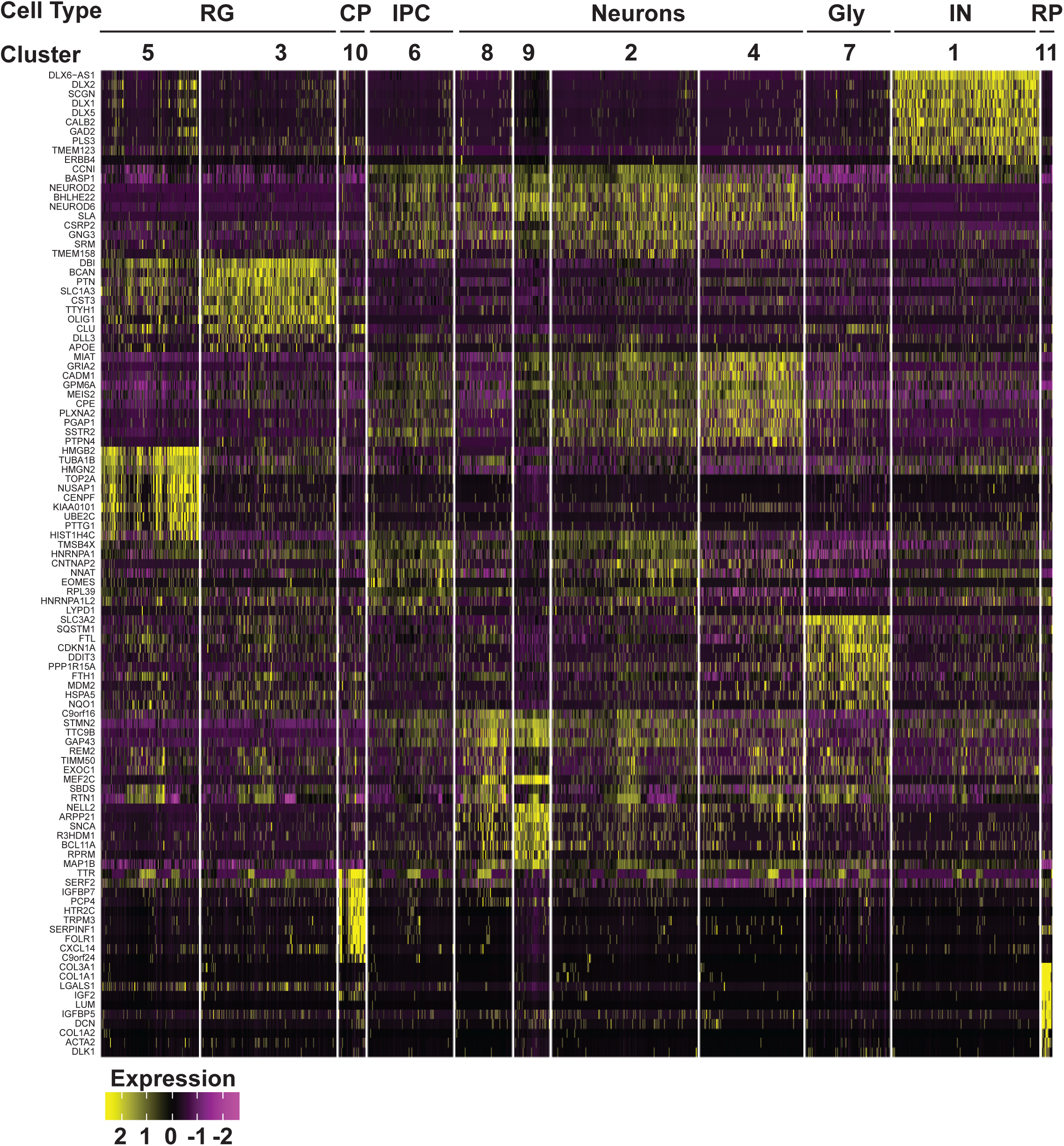
Gene expression heat map from 24 week old COs. The top 10 most differentially expressed genes per cluster are shown.

**Supplementary Figure 9.**
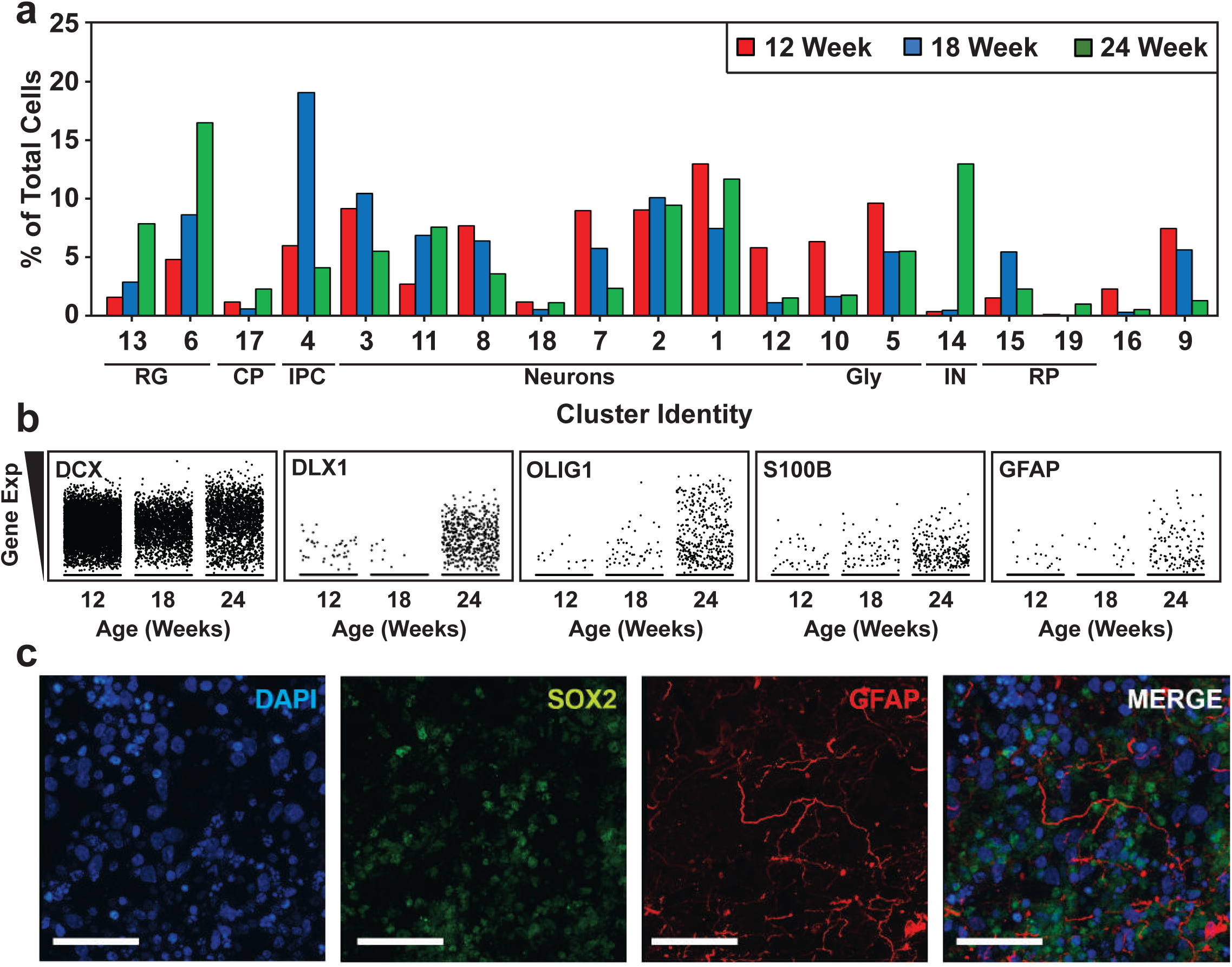
Cell type composition of hCOs changes with maturation. (**a**) Cluster frequency analysis depicting the percentage of cells at each timepoint that contributed to each cluster. RG; radial glial cells, oRG/astro; outer radial glial cells/astroglia, proRG; proliferative radial glial, CP; choroid plexus, IPC; intermediate progenitor cells, GLY; glycolytic signature, RP; Ribosomal Protein, IN; interneurons. (**b**) Scatterplots showing the expression of cell lineage markers including DCX (neurons), S100B (mature astrocytes), GFAP (astroglia/astrocytes), DLX1 (interneurons), and OLIG1 (oligodendrocyte precursors) at 12, 18 and 24 week time points. Note the general neuronal marker DCX, which is expressed similarly across the time points was used as a reference. (**c**) Astrocyte in 24 week old hCOs. The localization of GFAP (radial glia and astrocytes) and SOX2 (radial glia) in hCOs at 24 weeks of age, costained with DAPI was visualized by immunofluorescence microscopy. Scale Bar = 50 μm.

**Supplementary Figure 10.**
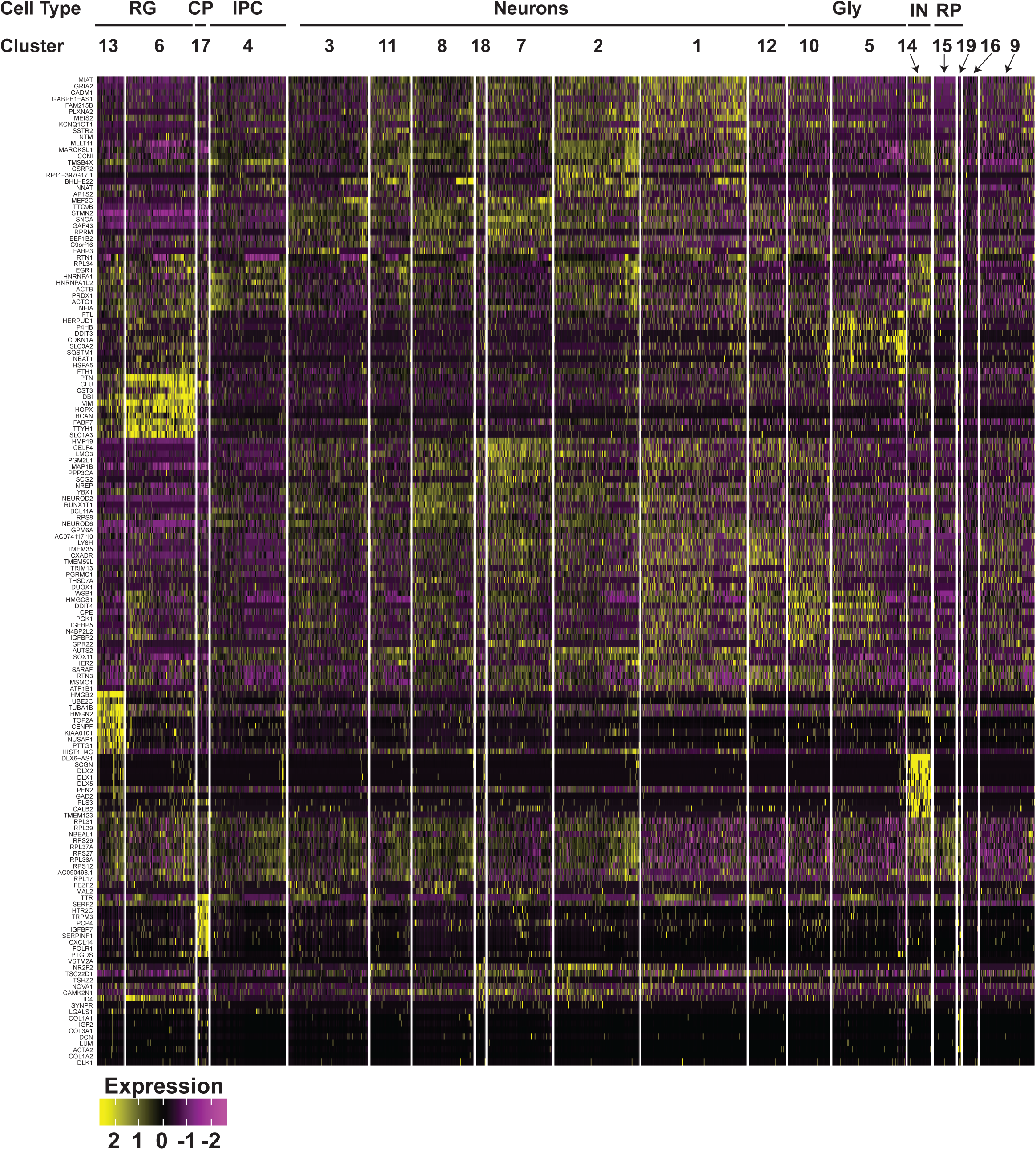
Gene expression heat map from combined 12, 18 and 24 week old COs. The top 10 most differentially expressed genes per cluster are shown.

**Supplementary Figure 11.**
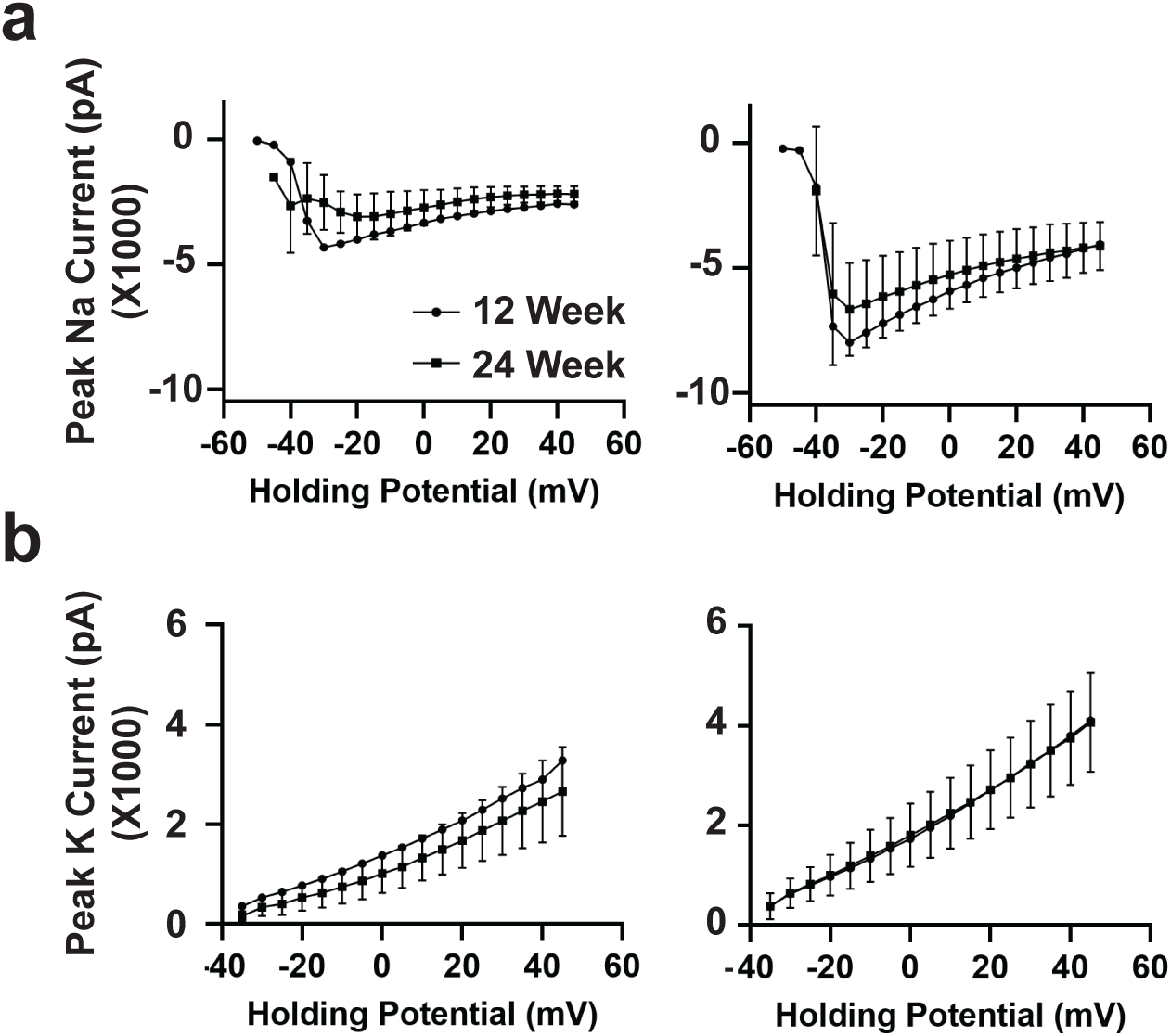
Electrophysiological analysis of 12 and 24 week old hCOs using whole cell patch clamping. (**a, b**) Na+ (**a**) and K+ (**b**) currents in developing and mature neurons analyzed in Figure 4 are plotted ± SEM.

**Supplementary Table 1:** Single-cell RNA sequencing (scRNA-seq) data analyses and statistics.

| **Time point**<br>**(weeks)** | **Batch** | **Sample Name** | **Total Raw Reads** | **Mapping rate to the mm10 transcriptome** | **Observed # of cells from Cellranger** | **Final # of Cells Analyzed post-processing** | **Total Genes post-processing** | **Mean Genes per cell post-processing** | **Mean UMIs per cell post-processing** |
| --- | --- | --- | --- | --- | --- | --- | --- | --- | --- |
| 12 | Batch 1 | CO1 | 311,904,172 | 44.90% | 2916 | 2413 | 17000 | 1994.58 | 4535.05 |
| 12 | Batch 1 | CO2 | 249,676,701 | 47.40% | 1698 | 1402 | 16092 | 1863.47 | 4349.66 |
| 12 | Batch 2 | CO3 | 305,610,493 | 49.60% | 1415 | 1196 | 15662 | 1717.15 | 4158.14 |
| 12 | Batch 2 | CO4 | 296,970,514 | 48.30% | 1856 | 1556 | 15992 | 1595.73 | 3496.43 |
| 12 | Batch 3 | CO5 | 299,860,802 | 46.40% | 1638 | 1367 | 15612 | 1582.65 | 3338.76 |
| 12 | Batch 3 | CO6 | 254,262,899 | 48.20% | 1462 | 1312 | 14494 | 1204.48 | 2377.26 |
| 18 | Batch 1 | CO1 | 231,219,972 | 39.70% | 3000 | 1302 | 14947 | 1139.10 | 2522.29 |
| 18 | Batch 2 | CO2 | 261,267,143 | 42.00% | 3000 | 746 | 14318 | 1400.24 | 3301.95 |
| 18 | Batch 3 | CO3 | 249,623,495 | 39.10% | 3000 | 942 | 14485 | 1136.37 | 2455.74 |
| 24 | Batch 1 | CO1 | 176,957,964 | 48.30% | 521 | 422 | 14005 | 1868.524 | 4649.917 |
| 24 | Batch 1 | CO2 | 226,313,736 | 47.00% | 514 | 436 | 13394 | 1688.03 | 4054.97 |
| 24 | Batch 1 | CO3 | 422,721,248 | 51.20% | 397 | 321 | 13058 | 1587.05 | 3866.52 |
| 24 | Batch 2 | CO4 | 224,554,922 | 55.20% | 264 | 217 | 12487 | 1656.78 | 4688.47 |
| 24 | Batch 2 | CO5 | 485,193,019 | 52.20% | 270 | 235 | 13495 | 2576.37 | 7381.49 |
| 24 | Batch 3 | CO6 | 97,086,115 | 50.80% | 270 | 212 | 12096 | 1775.474 | 4682.637 |
| 24 | Batch 3 | CO7 | 97,109,058 | 55.80% | 300 | 244 | 13476 | 2260.473 | 6701.987 |
| 24 | Batch 3 | CO8 | 442,525,699 | 49.90% | 604 | 519 | 14303 | 2074.306 | 5418.022 |

**Supplementary Table 2:**
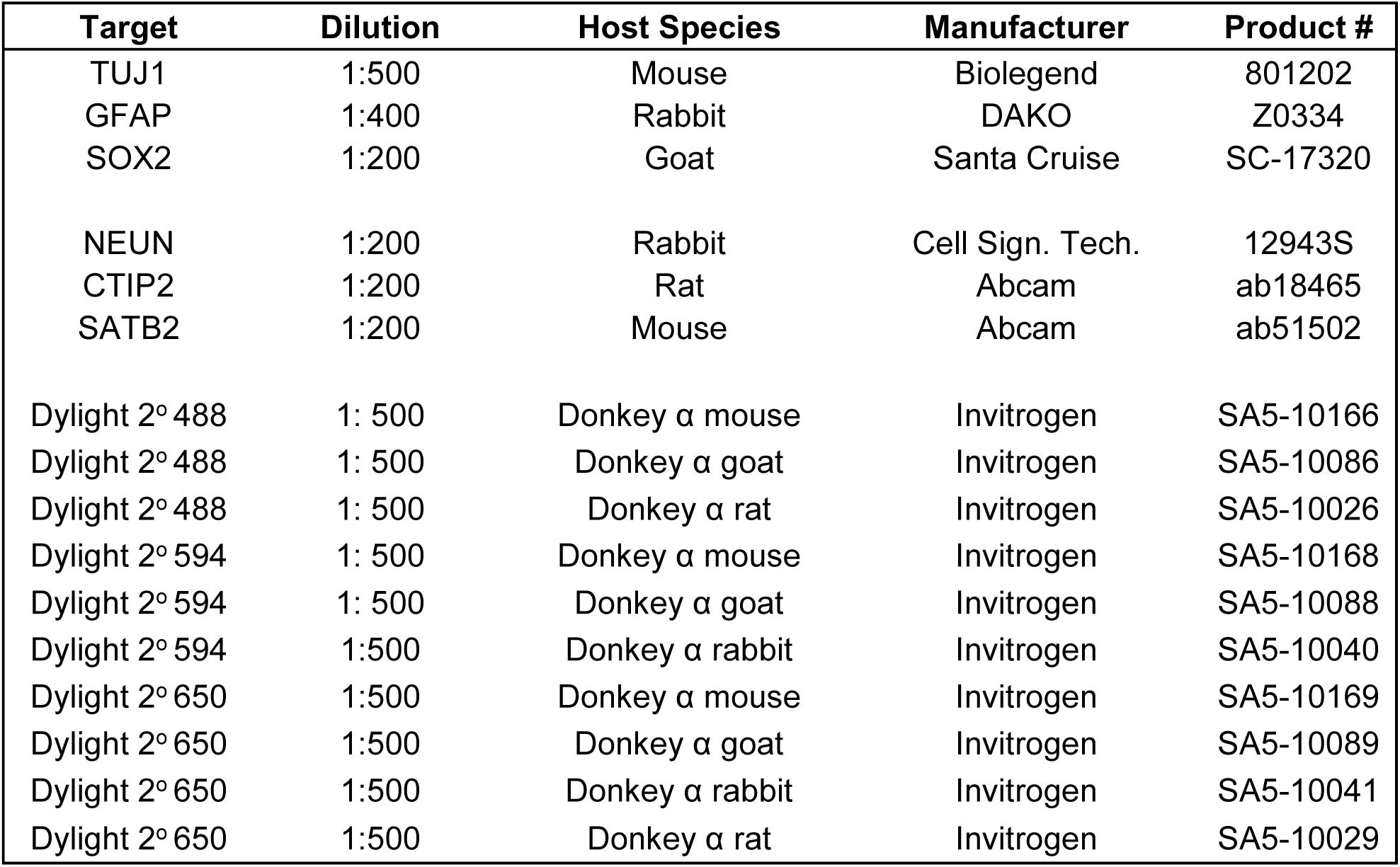
Antibody List.

